# Regional estimates of gross primary production applying the Process-Based Model 3D-CMCC-FEM vs. multiple Remote-Sensing datasets

**DOI:** 10.1101/2023.06.13.544712

**Authors:** D. Dalmonech, E. Vangi, M. Chiesi, G. Chirici, L. Fibbi, F. Giannetti, G. Marano, C. Massari, A. Nolè, J. Xiao, A. Collalti

**Affiliations:** Forest Modelling Laboratory, Institute for Agriculture and Forestry Systems in the Mediterranean, National Research Council of Italy (CNR-ISAFOM), 06128 Perugia, Italy; National Biodiversity Future Center (NBFC), 90133 Palermo, Italy; National Research Council of Italy, Institute of BioEconomy, (CNR-IBE), 50019 Sesto Fiorentino, Italy; GeoLab - Laboratory of Forest Geomatics, Dept. of Agriculture, Food, Environment and Forestry, Università degli Studi di Firenze, 50145 Firenze; Forest Ecology, Institute of Terrestrial Ecosystems, Department of Environmental Systems Science, ETH Zurich, Zurich, Switzerland; Research Institute for Geo-Hydrological Protection, National Research Council (CNR-IRPI), 06128 Perugia, Italy; School of Agricultural, Forest, Food and Environmental Sciences University of Basilicata, 85100 Potenza, Italy; Earth Systems Research Center, Institute for the Study of Earth, Oceans, and Space, University of New Hampshire, Durham, NH 03824, USA

**Keywords:** process-based forest model, wall-to-wall map, gross primary production, national forest inventory, Mediterranean region

## Abstract

Process-based Forest Models (PBFMs) offer the possibility to capture important spatial and temporal patterns of both carbon fluxes and stocks in forests, accounting for ecophysiological, climate and geographical variability. Yet, their predictive capacity should be demonstrated not only at the stand-level but also in the context of large spatial and temporal heterogeneity. For the first time, we apply a stand scale process-based model (3D-CMCC-FEM) in a spatially explicit manner at 1 km spatial resolution in a Mediterranean region in southern Italy. Specifically, we developed a methodology to initialize the model that comprehends the use of spatial information derived from the integration of remote sensing (RS) data, the national forest inventory data and regional forest maps to characterize structural features of the main forest species. Gross primary production (GPP) is simulated over the period 2005-2019 and the multiyear predictive capability of the model in simulating GPP is evaluated both aggregated as at species-level by means of independent multiple data sources based on different RS-based products. We show that the model is able to reproduce most of the spatial (∼2800 km^2^) and temporal (32 years in total) patterns of the observed GPP at both seasonal, annual and interannual time scales, even at the species-level. These new very promising results open the possibility of applying the 3D-CMCC-FEM confidently and robustly to investigate the forests’ behavior under climate and environmental variability over large areas across the highly variable ecological and bio- geographical heterogeneity of the Mediterranean region.

**Key Points:**

- We apply a process-based forest model on a regular grid at 1 km spatial resolution in a Mediterranean region.
- Initial forest state is estimated using spatially explicit input data derived from remote sensing and national forest inventory data.
- The 3D-CMCC-FEM shows comparably estimates in simulating both spatial and temporally the gross primary production, when compared to independent satellite-based products.

## 1 Introduction

Forest ecosystems absorb globally ∼2 Gigatonnes of Carbon (C) stocking the carbon in their biomass and soil, thus acting as a net carbon sink. In Europe alone, forest ecosystems, which cover about a 40%, currently act as a net carbon sink for ∼315 Megatonnes of CO_2_eq and compensate for about 8% of EU-27’s total greenhouse emissions (Verkerk et al., 2022). However, adverse climate impacts such as heat waves and drought (Allen et al., 2015; D’Andrea et al., 2020, 2021; Schuldt et al., 2020) and increasing natural disturbance rates (Grünig et al., 2023; Patacca et al., 2023) are all stressors which have potentially significant effects on current and future forest dynamics, jeopardizing the European forest ecosystems functioning and their carbon mitigation potential under future climate change (Schuldt et al., 2020; Senf et al., 2020; De Marco et al., 2022).

Nevertheless, ground data scarcity and short-term monitoring efforts still represent major challenges in studying the effects of climate change on forest dynamics in Mediterranean areas because are characterized by a large ecosystem heterogeneity and a biogeographically diverse structure shaped by human activity (Gauquelin et al., 2018; Médail et al., 2019; Peñuelas et al., 2017). As the Mediterranean region are known as a climate change ‘hotspot’ (Dubrovský et al., 2014; Noce et al., 2016), experiencing already increasing frequency in extreme events such heat waves and droughts (Vogel et al., 2021) it is thus crucial in these areas to provide large-scale forest monitoring and eventually predict the future state of forest ecosystems. Recent efforts have addressed the shortage of ancillary or ground data by integrating National Forest Inventories (NFI) data and high-resolution remote-sensing (RS) data. These initiatives produced comprehensive wall-to-wall maps of various forest variables, such as growing stocks volumes or biomass (Nord-Larsen & Schumacher, 2012; Waser et al., 2017; Chirici et al., 2020; Giannetti et al., 2022; Vangi et al., 2023). These maps represent meaningful data for and carbon cycle assessment (e.g. Vangi et al., 2023). In parallel, process-based forest models (PBFMs) are analytical tools developed and tested over a wide range of applications, because of their capability in simulating forest ecosystem even on long-term dynamics (Vacchiano et al., 2012; Bugmann & Seidl, 2022), carbon fluxes exchange and stocks (Chiesi et al., 2010; Reyer et al., 2014; Reyer, 2015; Dalmonech et al., 2022; Mahnken et al., 2022) under external environmental variability by accounting for population dynamics and inner physiological processes mechanistically (Pretzsch et al., 2008; Vacchiano et al., 2012; Maréchaux et al., 2021). On the other side, given the large amount of requested data for their initialization and parameterization, such models are mostly run at a very local scale, i.e. site level (one hectare or a bit more), where high-quality/measured ancillary data and meteorological data are available (Collalti et al., 2016; Suárez-Muñoz et al., 2023). Yet, initializing stand scale PBFMs from actual, measured forest state variables (or close to the observed states), rather than from equilibrium conditions, is the desirable option to implicitly take into account the climate and management history of the site, and to more realistically simulate the response of forests and their resilience, even in the context of climate change, and natural and anthropogenic disturbances (e.g. Pretzsch et al., 2008; Kannenberg et al., 2020; Zampieri et al., 2021).

Despite their undoubted utility, only recently, stand-scale PBFMs were applied on a regular grid (Sanchez-Ruiz et al., 2018; Minunno et al., 2019) with the purpose of estimating aggregated, country-level, carbon stocks and wood products. Yet, in southern Europe and in the Mediterranean the ability of a PBFMs to simulate the GPP at large scale is crucial, but largely overlooked, because photosynthesis respond sensitively to both meteorological and climate variability and spatial heterogeneity at the daily to decadal scales (e.g. Mahnken et al., 2022; Fernández-Martínez et al., 2023), therefore GPP can be, and has been, considered a good proxy of the ecosystem physiological functionality especially in Mediterranean forests (Collalti et al., 2018; Chen et al., 2023). The objective of this study is, thus, the application and the testing of the biochemical, biophysical, process-based forest model 3D-CMCC-FEM (Collalti et al., 2014, 2016, 2018), on a regular grid at 1 km spatial resolution in the Mediterranean area, initializing the model through the use of spatial information derived by integrating data of different nature: NFI data, RS-based wall-to-wall map, and regional forest maps to characterize structural features of the main forest species. The final aim is to simulate GPP at regional scale. As a case study, the model was tested over one of the southernmost regions in Italy, the Basilicata Region, which, as most regions in the Mediterranean basin, spans over a multitude of ecological, morphological and soil-type gradients and climate conditions. The capability of the model to simulate the GPP in terms of mean annual, seasonal and interannual variability is evaluated by comparing model results against a portfolio of different independent RS-based GPP-estimates. The 3D-CMCC-FEM GPP vs. RS-based GPP data agreement is presented and sources of uncertainties and challenges of applying a PBFMs with the presented modeling strategy are also discussed.

## 2. Materials and Methods

### 2.1 Study Area

The Basilicata region has a spatial extent of about 10,000 km^2^ and is located in southern Italy (Figure 1a). It is characterized by typical Mediterranean climate conditions with hot and dry summers and wet and mild winters. The region has been chosen to test the model in a complex biogeographic area with pronounced environmental gradients. The territory is characterized by about 47% of mountain areas represented by the Apennines Mountains, followed by hilly areas, about 45%, and then plain. Average annual precipitation varies between ∼500 and ∼2000 millimeters (mm) per year, mirroring the orographic complexity of the region and the proximities to the sea, with a west-to-east gradient from humid to dry sub-humid areas.

**Figure 1.**
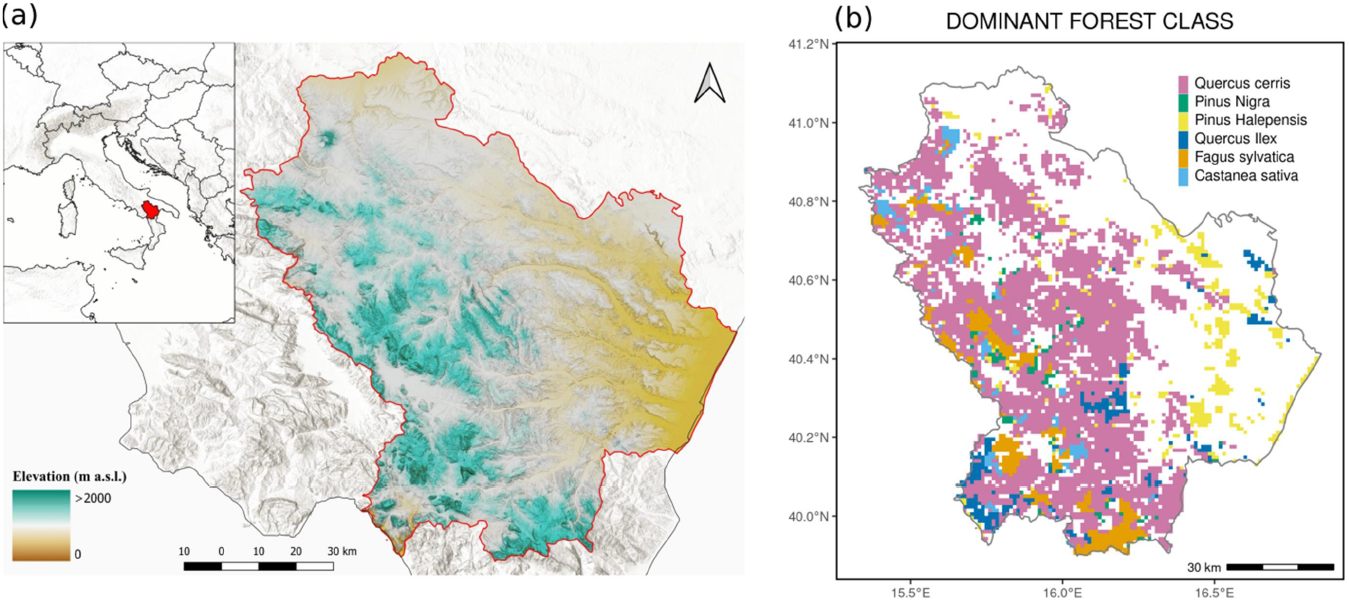
a) Study area in the Italian peninsula and elevation map of the Region Basilicata. The red line indicates the administrative limits of the region, b) distribution of the dominant forest class at 1×1 km spatial resolution.

According to the last NFI (INFC 2015), forest vegetation and other wooded lands occupy 392,412 hectares (ha), about 39% of the region. Deciduous species cover 54% of the forest area and are represented mainly by oaks spp (*Q. cerris* L., *Q. ilex* L.), which dominated the hilly areas between 400 and 1200 m above sea level (a.s.l.), and European beech (*Fagus sylvatica* L.) the main species above 1000 m a.s.l.. Coniferous species are less abundant and are represented mainly by pines spp (*P. halepensis* Mill., and *P. nigra* J.F. Arnold), often used for reforestation purposes (Figure 1b).

The study area was tessellated into a 1 km spatial resolution regional grid whose pixel area represent the best compromise between the forcing variables and the operative model resolution, for a total of ∼10,073 pixels. The regional grid served as a spatial reference grid for resampling the input data needed for the model initialization to 1 km spatial resolution.

### 2.2 The Process-based model 3D-CMCC-FEM

The 3D-CMCC-FEM (‘*Three Dimensional – Couple Model Carbon Cycle – Forest Ecosystem Module’*) is an ecophysiological, biogeochemical, biophysical process-based model which simulates the dynamic of carbon, water and nitrogen and the allocation through a cohort-structured forest stands (Collalti et al., 2014, 2016, 2017, 2018, 2020, 2022; Marconi et al., 2017; Dalmonech et al., 2022; Testolin et al., 2023), providing detailed output from daily to annual time scale of carbon fluxes and stocks. The model simulates forest growth and structural development at varying environmental conditions and different climate, atmospheric CO_2_ concentrations and forest management scenarios (Collalti et al., 2018; Dalmonech et al., 2022; Testolin et al., 2023).

The daily gross photosynthesis is simulated through the Farquhar–von Caemmerer–Berry biochemical model (Farquhar et al., 1980), modified for sun and shaded leaves (De Pury & Farquhar, 1997), and acclimated for temperature (Kattge & Knorr, 2007). The carbon and nitrogen allocation schemes are described extensively in Collalti et al. (2016, 2019, 2020) and Merganičová et al. (2019). Tree removal can occur via management (Testolin et al., 2023) or by natural mortality (Collalti et al., 2018). Self-thinning, age-related, carbon starvation, and background mortality represent the different types of mortalities simulated by the model (Collalti et al., 2016). Soil hydrology is simulated by means of a one-soil layer bucket model with free drainage. The plant water availability in the model is thus modulated by the soil depth (i.e. rooting depth until which the water is uptake to sustain leaf transpiration), because of the zero-dimensional soil model. Soil water stress operates on canopy exchange processes *via* stomatal and biochemical pathways (e.g. photosynthesis). An in-depth description of the model’s underlying characteristics, effects of climate change and model parameter sensitivity and uncertainty, as well as model limitations, is reported in Collalti et al. (2019).

### 2.3 Forest data source

The model requires the description of the forest structural characteristics: i.e. diameter at breast height (DBH), tree height (H), stand density (number of trees per cell) and age class, in order to be initialized and to run the simulations. In this work, to initialize the model for each grid cell of the matrix, we used data from the second NFI for 2005 (www.inventarioforestale.org). The NFI is based on a three-phase, systematic, unaligned sampling design with 1 km grid cells. In the first phase, 301,300 points were extracted and classified using aerial orthophotos into forest/non- forest categories. In the second phase, a field survey was carried out in a sub-sample of the first- phase points falling in the forest category, to collect qualitative information such as forest type, management, and property. Finally, in the third phase, for a sub-sample of 6782 points extracted from the second-phase points, a dendrometric survey was carried out for circular plots of a 13 m radius. All tree stems with DBH of at least 2.5 cm were callipered, and for a subsample, height was measured.

Field-survey data from the NFI were used to produce a ‘wall-to-wall’ map of the forest basal area at a 23 m spatial resolution. This map consists of random forests predictions of basal area per hectare for all 23 m spatial resolution forest pixels. The random forests model was trained using NFI plot-level data and Landsat and other RS-based datasets as predictors, including climate information such as minimum, mean, and maximum temperature, and daily precipitations from the E-OBS dataset, the land-only gridded daily observational dataset for Europe (see section 2.4). The statistical model fitting and tuning steps were carried out using the ‘*randomForest’* package in the statistical software R 4.0.5 (Liaw & Wiener, 2002). More information about the procedure can be found in Chirici et al. (2020), Vangi et al. (2021) and Giannetti et al. (2022). The pixel-level estimations of the basal area range between 5 and 43 m^2^ ha^−1^ with a mean value of 12 m^2^ ha^−1^, that is in line with the range reported in the context of NFI for the field plots data estimation.

The basal area data were then resampled to the regional grid at 1 km resolution. The resampled amp was then masked according to the regional forest map by Constantini et al. (2006). This map provided the forest type according to Barbati et al. (2007) and the development stage of the forests. Estimation of the forest structural data used to initialize the model in each regional grid cell, is described in section 2.5.1.

### 2.4 Forcing and soil data

The 3D-CMCC-FEM was forced with daily maximum (Tmax, °C) and minimum (Tmin, °C) air temperatures, precipitation (P, mm day^−1^), downward short-wave radiation at the surface (SW MJ m^−2^ day^−1^) and relative humidity (RH, %). Meteorological data for the period 2005-2019 were retrieved from the E-OBS v.23.1e gridded dataset (Cornes et al., 2018), which is provided at 0.1° decimal degree resolution The E-OBS dataset has already been used in environmental impact studies (e.g. Rita et al., 2020), climate scenarios bias correction (Dosio & Paruolo, 2011; Rojas et al., 2011) and benchmark activities (Moreno & Hasenauer, 2016; Herrera et al., 2019; Lorenz et al., 2019; Massari et al., 2020).

All the physical variables were bilinearly interpolated to the regional grid at 1 km resolution and the temperature data were corrected for the topographic effect by applying a lapse rate correction based on elevation differences between the E-OBS reference elevation and the finest and most accurate DEM (Digital Elevation Model) currently available in Italy, obtained in the framework of the TINITALY project. The TINITALYDEM is a national DEM of Italy at 10 m resolution (Tarquini et al., 2009) and for the lapse rate correction it was resampled at 1 km resolution of the regional grid. The lapse rate estimates of −5 °C km^−1^ for Tmax and −3 °C km^−1^ for Tmin, were derived from termo-pluviometric ground station measurements over an elevation transect in the Basilicata region.

The model was forced by global annual atmospheric CO_2_ concentrations from Dlugokencky and Tans (https://www.esrl.noaa.gov/gmd/ccgg/trends/), covering in total the years 2005-2019.

The model requires information on soil depth as well as soil texture for each grid cell. As a proxy of soil depth, we used the estimated depth available to root from the European Soil Database Derived Data product ESBD v2 (Hiederer, 2013) provided at 1 km resolution. The lower depth value of each class of the map was attributed, and a maximum of 1 m for the rooting depth was set. This boundary value can be considered a good approximation for European forests (Schenk & Jackson, 2005). Soil texture as a percentage of clay, silt, and sand was estimated from the pedological map of the region (year 2005).

The meteorological and geographic information were all re-projected using the same coordinate reference system WGS84/UTM zone 33 North (EPSG: 32633), and then resampled onto the regional grid at 1 km resolution. The main data analyses were performed using the computing language *R* (R Core Team 2021). Key packages used for data preprocessing included ‘*terra’* (Hijmans et al., 2022) and ‘*rgdal’* (Sumner & Hijmans, 2023)

### 2.5 Model simulations

#### 2.5.1 Model initialization

The 3D-CMCC-FEM model (v.5.6) was applied on the regional grid at 1 km spatial resolution to simulate the forest carbon dynamic starting in January 2005 until December 2019. The model requires the description of the forest attributes at the beginning of simulation in order to be initialized. The initial forest state was set according to a simplified model initialization of the forest aboveground structural complexity, i.e. for each grid cell, we determined the dominant forest species and estimated the average structural data: the average tree diameter at breast height (DBH), the average tree height (H), the stand density, and the average age class which represent the mandatory initial data for model initialization.

The following six key species were considered: European beech (*F. sylvatica* L.), Black pine (*P. nigra* J.F. Arnold), Sweet chestnut (*C. sativa* Mill.), Turkey oak (*Q. cerris* L.), Aleppo pine (*P. halepensis* Mill.) and Holm oak (*Q. ilex* L.) as representative of the most common forest types in the study area. The regional forest map was used to define for each 1 km grid cell the dominant forest species as the one covering the highest forest fraction (Figure 1b) and the average age class based on the development stage (provided by the regional forest map along with the forest class). Areas with dominance of maquis and other minor forest species (the latest accounts for ∼3.9% of the region and are not currently parameterized in the model) were masked out from the regional gird. The final dominant forest age classes, result in a total of ∼2800 km^2^, corresponding to ∼80 % of the entire regional forested area.

Tree density data of the NFI field plots were used to provide a representative estimate of the forest density in each grid cell. To do so, we zonally averaged the density data according to the regional grid and the dominant forest classes. The basal area map was then used in combination with the density map data to calculate an average cell-level DBH and to provide a 1 km resolution DBH map for each dominant species (data not shown).

To be consistent with the model processing and inherent logic, the H was calculated from the average DBH by applying the calibrated Chapmann-Richard equation (Richards, 1959), which links DBH and H, for each forest class with each cell. Starting from these mandatory structural variables, i.e. DBH and H, the model self–initializes the other state variables: i.e. leaf, stem, branches, coarse and fine root, reserves (non-structural carbon, NSC; which includes starch and sugars) carbon and nitrogen pools using a species-specific parameterization at the beginning of the model simulations. Species-specific model parameters (e.g. specific leaf area or maximum stomatal conductance, see Table S1) were retrieved and calibrated from literature data.

Specifically, the species-specific allometric equations linking DBH to H and DBH to stem biomass, were calibrated in this study Using the second NFI tree-level data.

#### 2.5.2 Simulations settings

Due to the relatively short time period covered by the simulations, i.e. 2005-2019, we did not consider any change in dominant species or land use. For the same reason, and because of the spatial resolution, a constant thinning rate implemented each year was considered as the only silvicultural intervention as similarly as in Gutsch et al. (2018). The thinning rate was an approximation derived from the forest management guidelines of the region as set to a yearly removal rate of 1%, corresponding to ∼20% of biomass removed in 20 years. Forests in protected areas of the Natura2000 network of the region are characterized by lower disturbance extension compared to the other forested areas according to the disturbance maps produced by Francini et al. (2021), and are assumed to be interested by a lower level of tree harvesting. For simplicity, we assume, thus, that mortality in protected area is mainly caused by natural and background mortality alone. Grid cells interested in fire events over the period 2005-2019, were identified using the national dataset of burnt areas from forest fires, produced by the Italian Forest Service (Comando Unità Forestali, Ambientali e Agroalimentari of Carabinieri). This dataset is acquired through a ground survey using Global Navigation Satellite System receiver (GNSS) and is available from 2005 to 2019. Grid cells where more than 30% of the forest area was interested in fire events were excluded from the analysis as the model does not simulate extended natural disturbances (i.e. fire and pests). This threshold was a compromise between excluding too many grid cells and including too many fire-disturbed areas.

### 2.6 Remote sensing - GPP datasets

Datasets based on RS-based data (or modeled by forcing with remote sensed data) are the most suitable candidate to assess the overall model capability at reproducing the GPP over large areas, due to their continuous spatial and temporal coverage. In order to make a more complete and comprehensive agreement assessment, we selected gridded GPP estimates from different independent sources as reference datasets, which are:

#### 2.6.1 GOSIF-GPP

The GOSIF GPP dataset (Li & Xiao, 2019b) is a recently developed GPP product that is based on the global OCO-2 based SIF product (Li & Xiao, 2019a). The GOSIF setup combines in a data-driven approach the remotely sensed sun-induced fluorescence (SIF), observed by the Orbiting Carbon Observatory-2 (OCO-2), the enhanced vegetation index (EVI) from the Moderate Resolution Imaging Spectroradiometer (MODIS) satellite data, meteorological data, i.e. photosynthetically active radiation (PAR), vapor pressure deficit (VPD), and air temperature obtained from the NASA reanalysis MERRA-2 data set to return a gridded SIF dataset (Li & Xiao, 2019a). Established relationships between the original OCO-2 SIF and flux tower GPP (Li et al., 2018; Xiao et al., 2019) were then used to provide the final gridded GPP product. For the model-data comparison in this study we used the monthly and annually aggregated ensemble mean of eight different GPP estimates resulting from different GPP-SIF relationships, (http://globalecology.unhedu). This GPP product, hereinafter simply referred as ‘GOSIF’, provides GPP estimates at 0.05 degree (corresponding to ∼5 km) spatial resolution aggregated on a monthly time step, over the years 2005-2019.

#### 2.6.2 CFIX-GPP

The CFIX dataset provides gridded GPP values covering Italy. The estimates are obtained combining meteorological and remotely sensed data within a Light Use Efficiency modeling approach (Veroustraete et al., 2002; Maselli et al., 2006). The original model version was further modified to simulate the GPP in water-limited, Mediterranean forest ecosystems by Maselli et al. (2009), who introduced a short-term water stress factor based on daily meteorological data. In particular, the authors utilized the 1-km normalized difference vegetation index (NDVI) from the Spot-VEGETATION imagery to linearly retrieve the fraction of absorbed photosynthetically active radiation (APAR), and the downscaled E-OBS meteorological data (both air temperature and precipitation)(Maselli et al., 2012; Fibbi et al., 2016) to retrieve the GPP of all Italian forests at 1-km spatial resolution. The accuracy of the product was assessed by comparison to Eddy-Covariance GPP data (Pastorello et al., 2020) collected at several EC sites spread over the Italian peninsula and for different forest types, showing satisfactory performances (Chirici et al., 2016). The currently utilized product, provides forest GPP estimates at 1 km spatial resolution aggregated on a monthly time step, over the years 2005-2013.

#### 2.6.3 FLUXCOM-GPP

The FLUXCOM dataset is an upscaled product derived from different Machine Learning-based approaches (ML) combining data from the FLUXNET network of eddy covariance towers with RS and meteorological data as predictors (Jung et al., 2019; Pastorello et al., 2020). In this study we used the FLUXCOM-RS dataset which embeds in its statistical processing, RS land products at 8-day resolution from the MODIS instrument, such as EVI, fAPAR, and the land surface temperature (see Jung et al. (2019) for a thorough description of the FLUXCOM products). Compared to other products of the FLUXCOM-database, the FLUXCOM-RS dataset has a finer spatial resolution, 0.083 degree (corresponding to ∼8 km) spatial resolution, allowing to better deal with the complex topography of the region under study.

The FLUXCOM-RS dataset, hereinafter simply referred to as FLUXCOM, has been widely used as a reference dataset in several inter-comparison studies with both model and other reference datasets (O’Sullivan et al., 2020; Wang et al., 2021; Zhang & Ye, 2021). The selected dataset is provided as the ensemble mean of monthly data computed from multiple ML algorithms covering the period 2000-2015 (www.fluxcom.org).

### 2.7 3D-CMCC-FEM GPP versus RS-based GPP

For comparability with the 3D-CMCC-FEM GPP results, all the RS-based datasets were resampled to the regional grid at 1 km resolution and reprojected using the WGS84/ UTM zone 33 North coordinate reference system (EPSG: 326633) and compared at both spatial and temporal level against the 3D-CMCC-FEM GPP data. Results at species-level are first shown as aggregated data over regional level and then shown at species-level.

#### 2.7.1 Spatial variability analyses

The spatial agreement between the 3D-CMCC-FEM GPP and the RS-based GPP data was assessed by considering: Root Mean Square Error (RMSE), Relative Difference (RD, expressed as: (modelled – observed) / observed * 100, where ‘observed’ stands for RS-based data and ‘modeled’ for 3D-CMCC-FEM GPP), Pearson’s correlation (r), and the SPAtial EFficiency (SPAEF) metrics. The SPAEF (see Eq. 1) is an integrated evaluation index which considers the Pearson’s spatial correlation (*r*), the fraction of the coefficient of variation *β* = (*σ*_sim_/*μ*_sim_)/(*σ*_obs_/*μ*_obs_) and the model-data histogram intersection 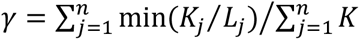, where *n* is the number of bins in the histogram and K and L are the histogram for observed and simulated data (Koch et al., 2018). SPAEF is equal to 1 in the case of a perfect match and 0 in case of mismatch.

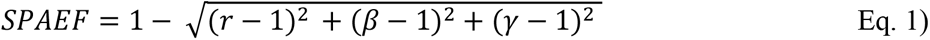

The statistics were computed on the mean annual GPP and seasonal GPP: i.e. winter (December, January, February: DJF), spring (March, April, May: MAM), summer (June, July, August: JJA) and autumn (September, October, November: SON) for 3D-CMCC-FEM and RS-based data.

#### 2.7.2 Temporal variability analyses

The temporal agreement between the 3D-CMCC-FEM GPP and the RS-based GPP data was assessed by considering RMSE, RD, the Fractional Variance (FV), which returns values bounded between −2 and 2 and it is equal to 0 when ‘modelled’ and ‘observed’ have the same variance (Janssen & Heuberger, 1995). The Pearson’s correlation coefficient *r* was also used at grid-cell level, which returns the direction of the correlation i.e. model-data correspondence of the sign of year to year variability. Conversely to the spatial analysis the mean seasonal cycle (MSC) was also considered here. Anomalies were calculated for the model and data by first removing the long-term linear trend, and normalized by their standard deviation.

Both the 3D-CMCC-FEM and the RS-based GPP climate sensitivity in the summer period has been evaluated through the Standardized Precipitation Evaporation Index, SPEI (Vicente-Serrano et al., 2010). The SPEI drought index helps to highlight periods of wetter or drier conditions. This is a multi-scalar meteorological drought index based on a statistical transformation of the climatic water balance, i.e. precipitation *minus* potential evapotranspiration. We computed the SPEI using the simplistic Hargreaves equations for the potential evapotranspiration calculation and considered different time scales, with the aim to cover the growing season period before the month of August. Following a similar approach as in Mahnken et al. (2022), we regressed the residuals of the 3D-CMCC-FEM and RS-based datasets anomalies against the SPEI values as the predictor, and the slope of the linear regression computed. Values of the slope close to 0 indicate that the 3D-CMCC-FEM shows a GPP-sensitivity to SPEI like the RS-based dataset.

The overall analyses where carried out considering the entire RS-based dataset available years, thus 2005-2013 for CFIX, 2005-2015 for FLUXCOM and 2005-2019 for GOSIF (i.e. 32 years in total).

## 3 Results

### 3.1 Spatial variability analysis

The 3D-CMCC-FEM simulated average annual GPP for the period 2005-2019 are shown in Figure 2, with overall values ranging from ∼600 up to ∼2200 gC m^−2^ yr^−1^. 3D-CMCC-FEM GPP follows a west-east gradient with higher productivity over the western side of the region in correspondence with the more humid and the more productive beech-dominated areas; the forested areas over the plain zones are mainly dominated by Mediterranean pine species, which show the lowest GPP values (Figure 2). At annual level (multiyear mean annual) better correlations between 3D-CMCC-FEM and RS-based GPP data are with GOSIF and FLUXCOM (*r* = 0.77 and 0.67, respectively; Table 1) as also for SPAEF metric (SPAEF = 0.62 and 0.59, respectively) while lower RMSE and RD are when 3D-CMCC-FEM GPP is compared with GOSIF and CFIX (RMSE = 235.8 and 221.9 gC m^−2^ yr^−1^, and RD = −4.3 and 0.01%, respectively). At seasonal level, 3D-CMCC-FEM GPP better correlates with GOSIF and CIFX during summer (RMSE = 181.3 and 123.8 g C m^−2^ yr^−1^, RD = −13.7 and 10.7% and *r* = 0.79 and 0.8, respectively). Similarly, also in spring 3D-CMCC-FEM better correlates, although with lower values, with GOSIF and CIFX (RMSE = 104.4 and 103.6 gC m^−2^ yr^−1^, RD = 9 and −7.9% and *r* = 0.4 and 0.28, respectively). In winter 3D-CMCC-FEM GPP better correlates with CIFX and FLUXCOM (*r* = 0.55 and 0.5, and SPAEF = −1.26 and 0.53, respectively) but with slightly lower RMSE values when compared with GOSIF and FLUXCOM (RMSE = 54.3 and 39.97 gC m^−2^ yr^−1^). During autumn 3D-CMCC-FEM is in agreement with GOSIF and CIFX (RMSE = 64.47 and 52.41 gC m^−2^ yr^−1^, RD = 2.4 and 15% and *r* = 0.74 and 0.67, respectively).

**Figure 2.**
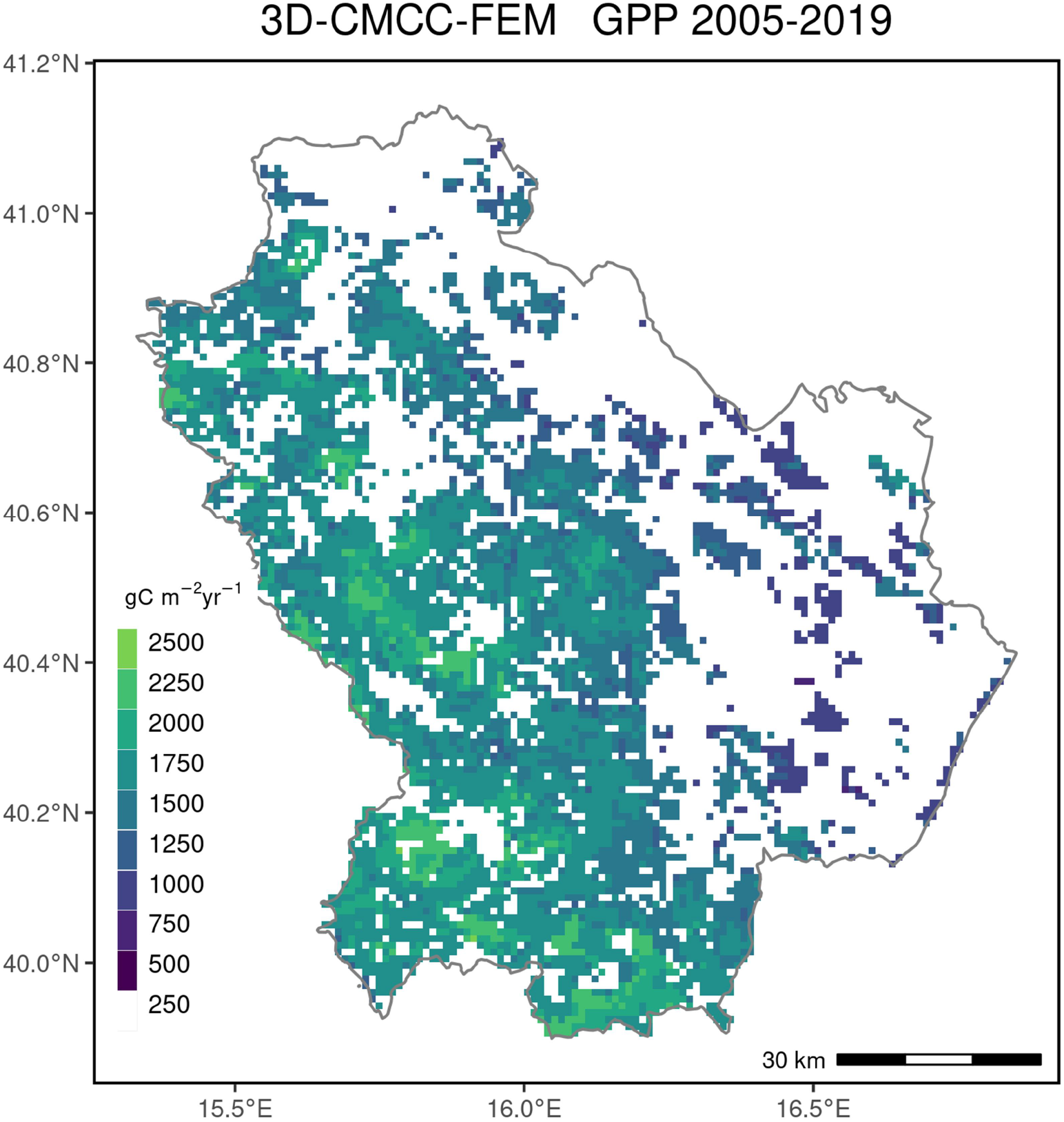
3D-CMCC-FEM mean annual GPP values (gC m^−2^ yr^−1^) for the period 2005-2019 at 1 km spatial resolution. White areas indicate areas not simulated by the 3D-CMCC-FEM.

**Table 1.**
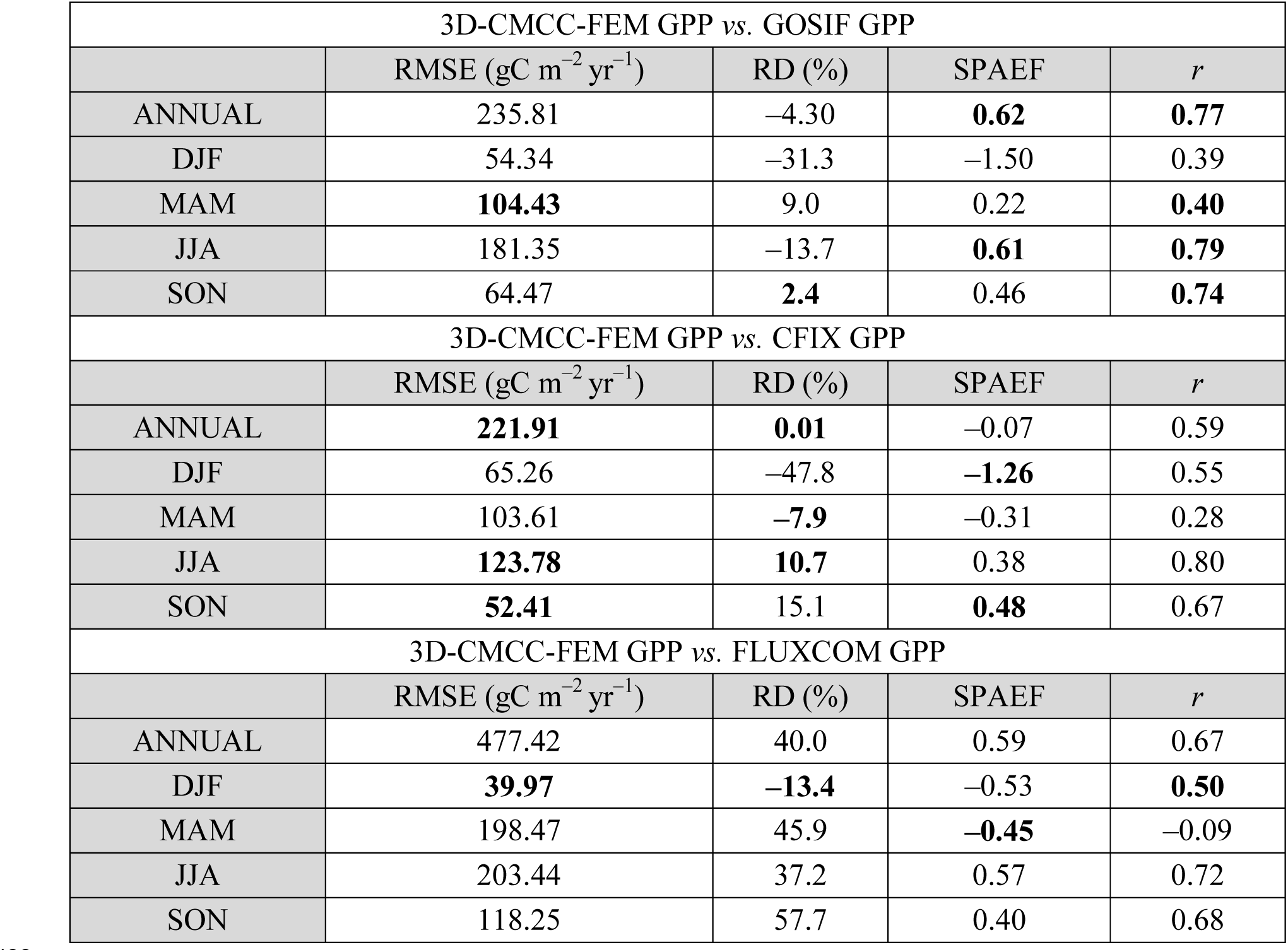
Spatial comparison of the 3D-CMCC-FEM GPP vs. RS–based GPP. RMSE= root mean square error (gC m^−2^ yr^−1^), RD = relative difference (%), SPAEF = SPatial EFficiency, *r* = Pearson’s correlation. DJF, winter months; MAM, spring months, JJA, summer months; SON, autumn months. In bold values with the best agreement between 3D-CMCC-FEM GPP and RS-based GPP.

### 3.2 Temporal variability analysis

Generally, the 3D-CMCC-FEM GPP shows overall lower correlations of the temporal (which corresponds to the interannual variability, IAV) vs. spatial comparison for the mean annual GPP. Similarly, as in the spatial analysis, 3D-CMCC-FEM GPP shows better agreement with GOSIF (RMSE = 231.6 gC m^−2^ yr^−1^, RD = −4.6% and *r* = 0.4, see Table 2 for overall statistics and Figure 3, 4, 5, 6 for the maps) and the lowest Fractional Variance between the RS-based dataset considered (FV = 0.83, see also Standard Deviation analysis Figure S1). At seasonal level, during summer, 3D-CMCC-FEM GPP correlates better with the FLUXCOM (*r* = 0.62 and FV = 1.64) although with the highest RMSE (182.53 gC m^−2^ yr^−1^) among the RS-based dataset considered. Conversely, in spring 3D-CMCC-FEM GPP correlates better with CFIX (*r* = 0.31) and with the lowest RMSE (86.81 gC m^−2^ yr^−1^) and RD (–7.9%). In winter modeled GPP values from 3D-CMCC-FEM better correlates with CFIX (*r* = 0.69) but with lower RMSE, RD and FV values when compared with FLUXCOM (RMSE = 36.24 gC m^−2^ yr^−1^, RD = −24.1 and FV = 1.77). Also during autumn season, the 3D-CMCC-FEM GPP better correlates with CFIX (*r* = 0.3) and with the lowest RMSE (57.97 gC m^−2^ yr^−1^) although the lowest Relative Difference is when compared with GOSIF (RD = 1.7%).

**Figure 3.**
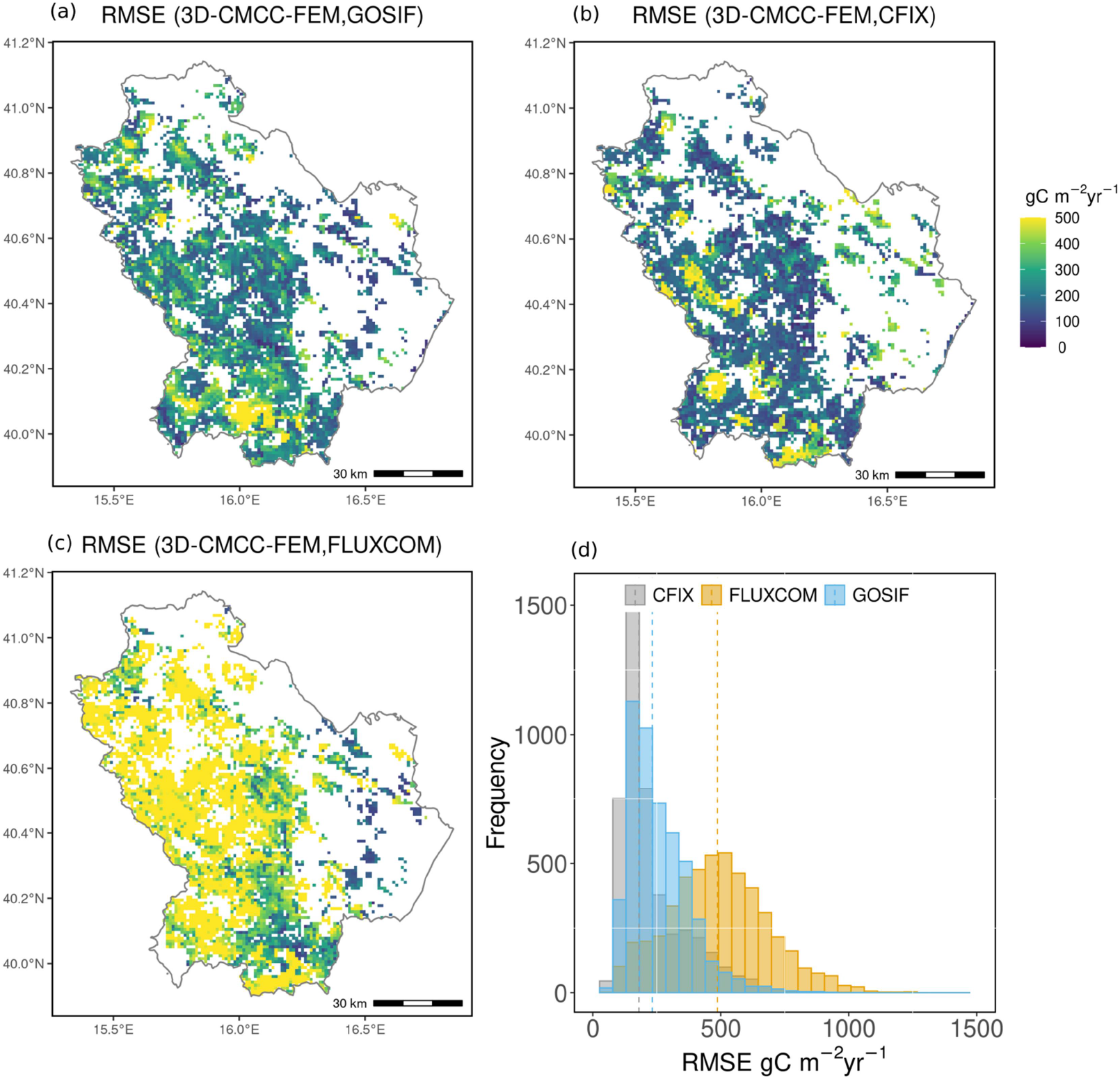
Root Mean Square Error (RMSE, gC m^−2^ yr^−1^) between mean annual 3D-CMCC-FEM GPP (gC m^−2^ yr^−1^) and the RS-based GPP: a) 3D-CMCC-FEM GPP vs. GOSIF GPP for the period 2005-2019, b) 3D- CMCC-FEM GPP vs. CFIX GPP for the period 2005-2013, c) 3D-CMCC-FEM GPP vs. FLUXCOM GPP for the period 2005-2015; d) histogram of the relative difference, dashed lines indicate the median value. White areas on the maps indicate areas not simulated by the 3D-CMCC-FEM.

**Figure 4.**
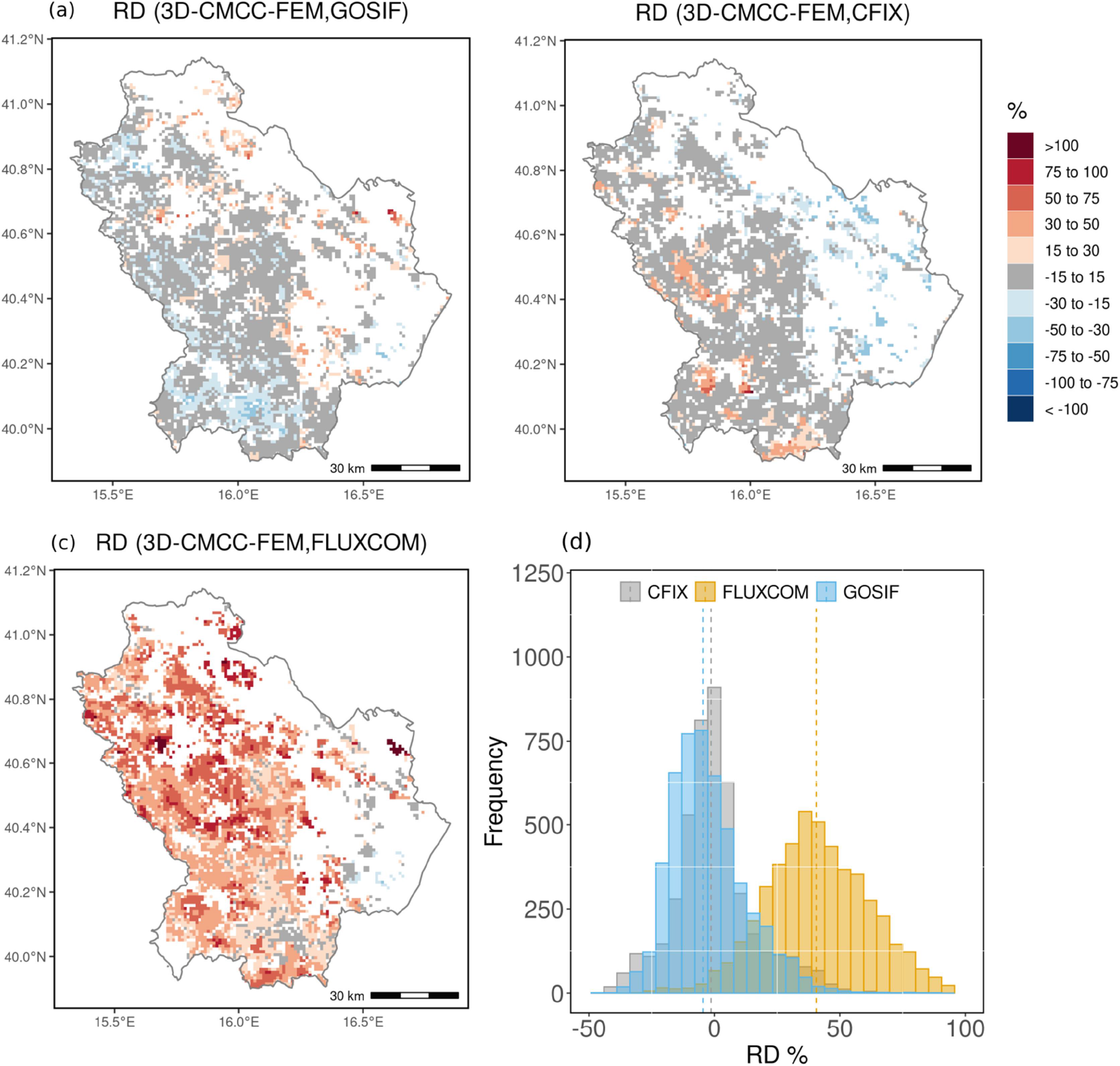
Relative Difference (RD, %) between mean annual 3D-CMCC-FEM GPP and the RS-based GPP: a) 3D-CMCC-FEM GPP vs. GOSIF GPP for the period 2005-2019, b) 3D-CMCC-FEM GPP vs. CFIX GPP for the period 2005-2013, c) 3D-CMCC-FEM GPP vs. FLUXCOM GPP for the period 2005-2015; d) histogram of the relative difference, dashed lines indicate the median value. White areas on the maps indicate areas not simulated by the 3D-CMCC-FEM.

**Figure 5.**
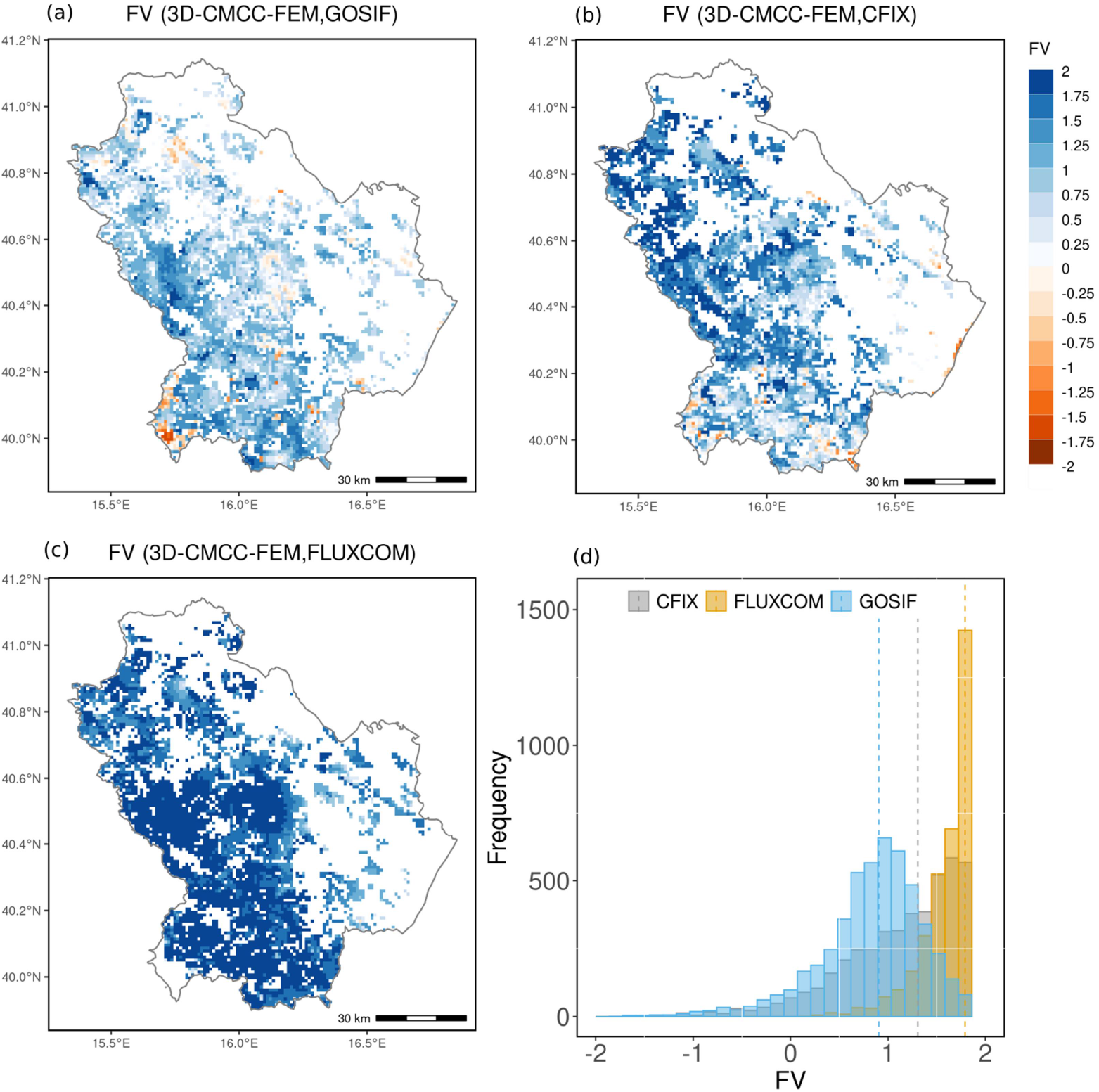
Fractional Variance (FV) between mean annual 3D-CMCC-FEM GPP and the RS-based GPP: a) 3D-CMCC-FEM GPP vs. GOSIF GPP for the period 2005-2019, b) 3D-CMCC-FEM GPP vs. CFIX GPP for the period 2005-2013, c) 3D-CMCC-FEM GPP vs. FLUXCOM GPP for the period 2005-2015; d) histogram of the relative difference, dashed lines indicate the median value. White areas on the maps indicate areas not simulated by the 3D-CMCC-FEM.

**Figure 6.**
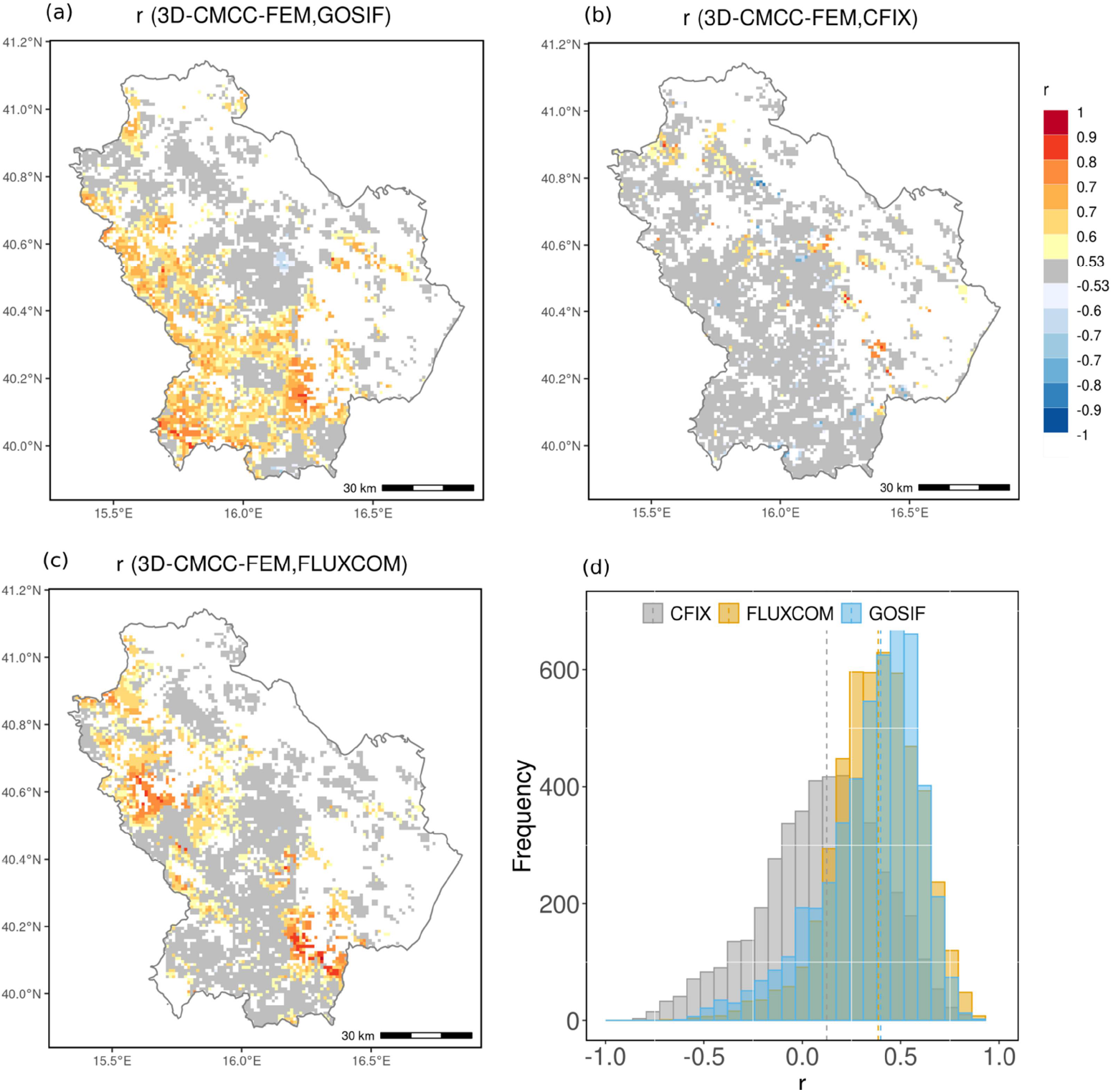
Pearson’s correlation (*r*) between mean annual 3D-CMCC-FEM GPP and the RS-based GPP: a) 3D- CMCC-FEM GPP vs. GOSIF GPP for the period 2005-2019, b) 3D-CMCC-FEM GPP vs. CFIX GPP for the period 2005-2013, c) 3D-CMCC-FEM GPP vs. FLUXCOM GPP for the period 2005-2015; d) histogram of the relative difference, dashed lines indicate the median value. Grey areas on the maps indicate where correlations are not significant. White areas on the maps indicate areas not simulated by the 3D-CMCC- FEM.

**Table 2.** Temporal comparison of the 3D-CMCC-FEM GPP vs. RS–based GPP. RMSE= root mean square error (gC m^−2^ yr^−1^); RD = relative difference (%); FV = fractional variance; *r* = Pearson’s correlation. Metrics are first computed at grid cell level and reported as the median value. DJF, winter months; MAM, spring months, JJA, summer months; SON, autumn months; MSC, mean seasonal cycle. In bold values with the best agreement between 3D-CMCC-FEM GPP and RS-based GPP.

| 3D-CMCC-FEM GPP vs. GOSIF GPP |  |  |  |  |
| --- | --- | --- | --- | --- |
|  | RMSE (gC m<sup>-2</sup> yr<sup>-1</sup>) | RD (%) | FV | *r* |
| ANNUAL | 231.60 | -4.6 | **0.83** | **0.40** |
| DJF | 54.64 | -40.6 | 1.70 | 0.51 |
| MAM | 105.61 | 10.4 | **0.77** | 0.15 |
| JJA | 146.72 | -12.7 | 0.97 | 0.53 |
| SON | 77.33 | **1.7** | **0.38** | 0.17 |
| MSC | / | / | / | 0.91 |
| 3D-CMCC-FEM GPP vs. CFIX GPP |  |  |  |  |
|  | RMSE (gC m<sup>-2</sup> yr<sup>-1</sup>) | RD (%) | FV | *r* |
| ANNUAL | **179.55** | **-1.3** | 1.17 | 0.12 |
| DJF | 66.67 | -58.7 | **1.09** | **0.69** |
| MAM | **86.81** | **-7.9** | 1.36 | **0.31** |
| JJA | **86.12** | **7.5** | **0.68** | **0.65** |
| SON | **57.97** | 15.2 | 0.65 | **0.30** |
| MSC | / | / | / | **0.95** |
| 3D-CMCC-FEM GPP vs. FLUXCOM GPP |  |  |  |  |
|  | RMSE (gC m<sup>-2</sup> yr<sup>-1</sup>) | RD (%) | FV | *r* |
| ANNUAL | 486.65 | 40.6 | 1.69 | 0.39 |
| DJF | **36.24** | **-24.1** | 1.77 | 0.55 |
| MAM | 209.14 | 49.4 | 1.66 | 0.26 |
| JJA | 182.53 | 37.9 | 1.64 | 0.62 |
| SON | 120.33 | 54.7 | 1.41 | 0.17 |
| MSC | / | / | / | 0.92 |

At the Mean Seasonal Cycle level (thus comparing month per month average GPP values) 3D-CMCC-FEM GPP well correlates with all RS-based dataset with *r* varying from 0.95 (when compared with CFIX) to 0.91 (when compared with GOSIF)(see Figure 7, Table 2 and Figure S2)

**Figure 7.**
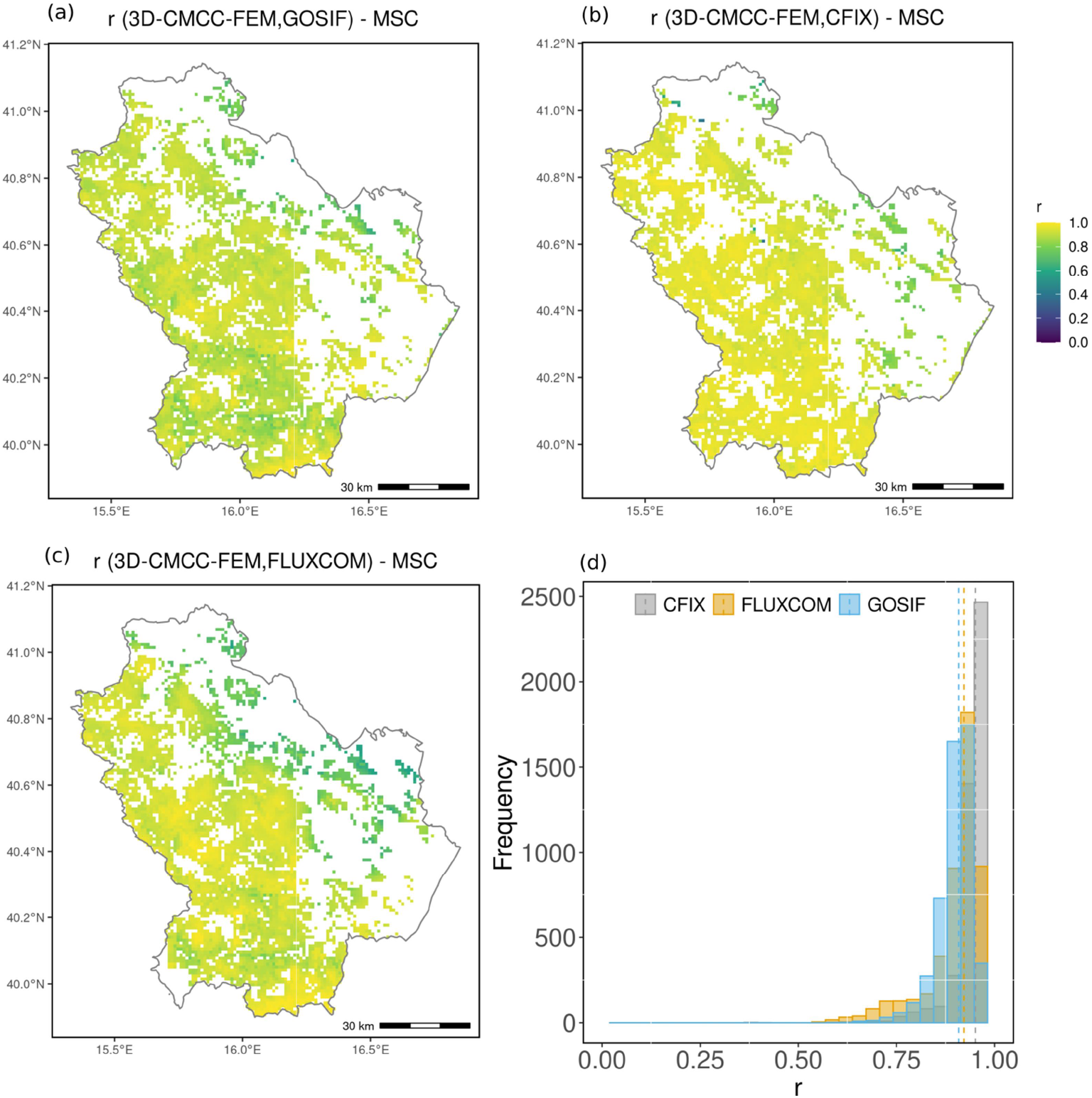
Pearson’s correlation (*r*) between Mean Seasonal Cycle (MSC) 3D-CMCC-FEM GPP and the RS- based GPP: a) 3D-CMCC-FEM GPP vs. GOSIF GPP for the period 2005-2019, b) 3D-CMCC-FEM GPP vs. CFIX GPP for the period 2005-2013, c) 3D-CMCC-FEM GPP vs. FLUXCOM GPP for the period 2005-2015, d) histogram of the r, dashed lines indicate the median value. White areas on the maps indicate areas not simulated by the 3D-CMCC-FEM.

### 3.3 SPEI

We first computed the SPEI at a time-scale of 5 months (SPEI5) representing the spring and summer period where the correlation between the RS-based summer GPP and SPEI were stronger. We estimate whether the 3D-CMCC-FEM data and RS-based data differ in GPP sensitivity toward interannual variation in SPEI by computing the slope of the regression between 3D-CMCC-FEM vs. RS-based differences of GPP and the SPEI. Generally, results show how, when considering CFIX and FLUXCOM, the slope is not significantly different from 0 for almost the entire simulation domain (Figure 8). Such a behavior indicates that 3D-CMCC-FEM and RS-based GPP summer anomalies respond to SPEI5 similarly. When comparing with the GOSIF the slope is still not significant over large areas. However, in clustered areas, corresponding to about a 20% of the simulation domain, the slope is positive and significant showing that the 3D-CMCC-FEM response to aridity is stronger than the GOSIF data (Figure 8).

**Figure 8.**
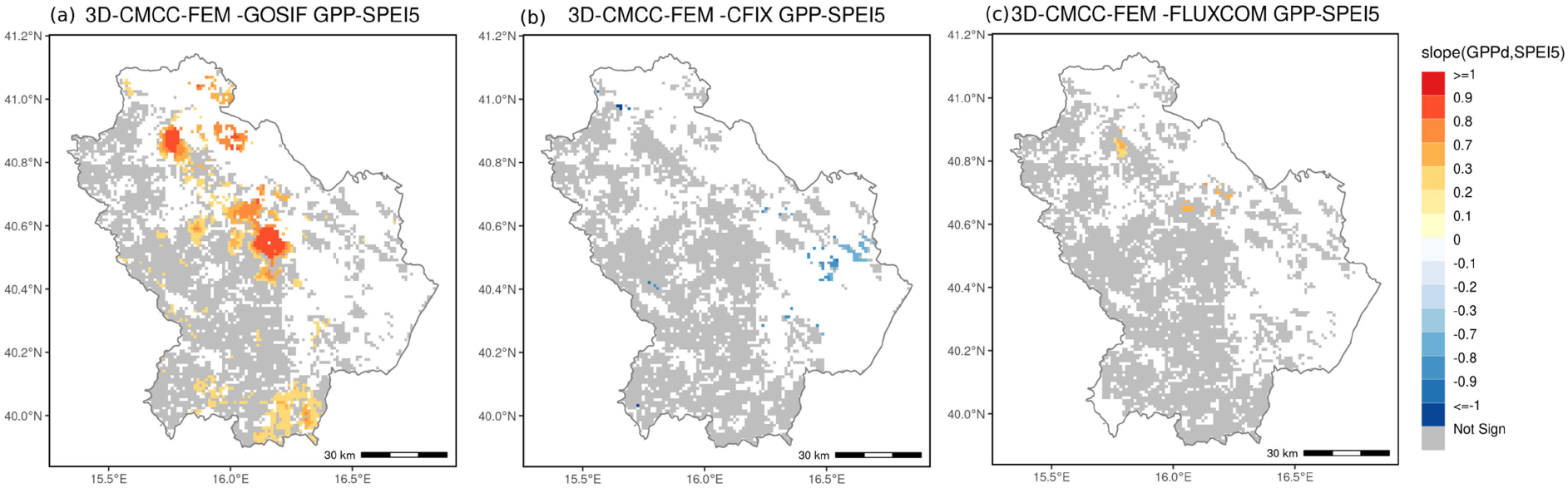
Spatial distribution of the slope of the linear regression of the summer GPP residuals and SPEI5 for: a) 3D-CMCC-FEM vs. GOSIF, b) 3D-CMCC-FEM vs. CFIX, c) 3D-CMCC-FEM vs. FLUXCOM. Grey areas indicate the slope is not significantly different from 0 at p<0.05. White areas on the maps indicate areas not simulated by the 3D-CMCC-FEM.

### 3.4 Species-level comparison

Data analysis at species level reveals as 3D-CMCC-FEM model GPP tends to correlate better for some species for some RS-based datasets than other (Table 3 and Figure 9). 3D-CMCC-FEM GPP correlates well for *Q. cerris* and *Q. ilex* with GOSIF (*r* = 0.73 and 0.82) but with low RMSE and RD for CFIX (RMSE = 141.09 and 141.99 gC m^−2^ yr^−1^, and RD = −2.69 and 3.39%, respectively). In all cases 3D-CMCC-FEM GPP for *F. sylvatica* are slightly far from results from all RS-based datasets with better correlations with CFIX (*r* = 0.43) and lower RMSE and RD values for GOSIF (193.35 gC m^−2^ yr^−1^ and 3.43%). Satisfactorily correlations are shown for *C. sativa* with the GPP values of FLUXCOM (*r* = 0.61) but with lower RMSE and RD with GOSIF (200.45 gC m^−2^ yr^−1^ and −1.71%). For Pinus species (both as *P. halpensis* and *P. nigra*), 3D- CMCC-FEM GPP shows to better correlates and with lower RMSE and RD values with GOSIF (*r* = 0.84, RMSE = 170.84 gC m^−2^ yr^−1^ and RD = −7.91% for *P. halepensis*; and *r* = 0.65, RMSE = 223.92 gC m^−2^ yr^−1^ and RD = 7.85% for *P. nigra*) than with CFIX and FLUXCOM (Table 3 and Figure 9).

**Figure 9.**
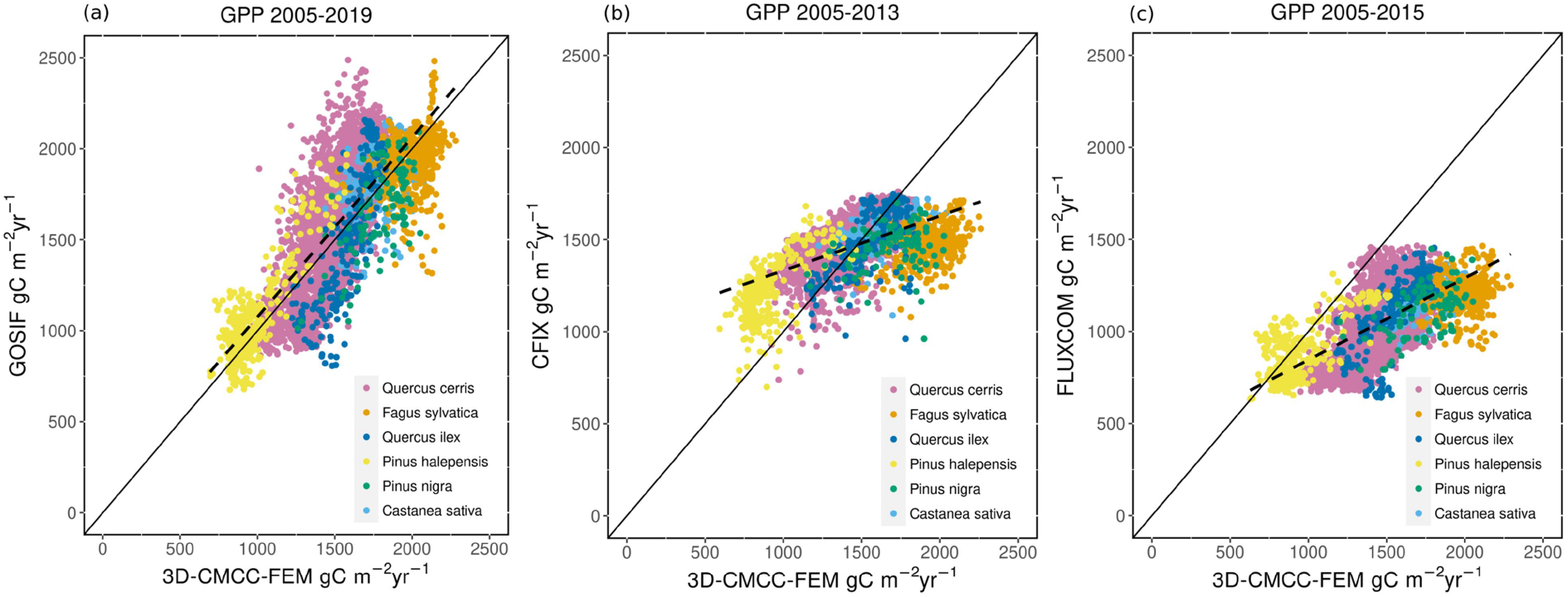
Mean annual 3D-CMCC-FEM GPP vs. RS-based GPP scatter plot (gC m^−2^ yr^−1^) at the species-level: a) the 3D-CMCC-FEM GPP vs. GOSIF b) 3D-CMCC-FEM GPP vs. CFIX and c) 3D-CMCC-FEM GPP vs. FLUXCOM estimates (black line is the 1:1 line, dashed line is the linear fit). Each point represents data at grid cell level, different colors indicate the different species considered.

**Table 3.** Comparison of the 3D-CMCC-FEM GPP vs. RS-based GPP at single species level. RMSE= root mean square error (gC m^−2^ yr^−1^), RD = relative differences (%), r = Pearson’s correlation. Metrics are first computed at grid cell level and reported as the median value. In bold values with the best agreement between 3D-CMCC-FEM GPP and RS-based GPP.

| 3D-CMCC-FEM GPP vs. GOSIF GPP |  |  |  |
| --- | --- | --- | --- |
|  | RMSE (gC m<sup>-2</sup> yr<sup>-1</sup>) | RD (%) | *r* |
| *Quercus cerris* | 245.66 | -6.33 | **0.73** |
| *Fagus sylvatica* | **193.35** | **3.43** | 0.28 |
| *Quercus ilex* | 267.65 | 3.96 | **0.83** |
| *Pinus halepensis* | 170.84 | -7.91 | **0.85** |
| *Pinus nigra* | **223.92** | **7.86** | **0.66** |
| *Castanea sativa* | **200.45** | **-1.72** | 0.25 |
| 3D-CMCC-FEM GPP vs. CFIX GPP |  |  |  |
|  | RMSE (gC m<sup>-2</sup> yr<sup>-1</sup>) | RD (%) | *r* |
| *Quercus cerris* | **141.1** | **-2.7** | 0.60 |
| *Fagus sylvatica* | 477.4 | 31.1 | **0.43** |
| *Quercus ilex* | **141.9** | **3.4** | 0.70 |
| *Pinus halepensis* | 355.1 (28%) | -26.8 | 0.65 |
| *Pinus nigra* | 314.5 (21%) | 16.9 | 0.25 |
| *Castanea sativa* | 203.3 | 7.0 | 0.36 |
| 3D-CMCC-FEM GPP vs. FLUXCOM GPP |  |  |  |
|  | RMSE (gC m<sup>-2</sup> yr<sup>-1</sup>) | RD (%) | *r* |
| *Quercus cerris* | 440.4 | 39.1 | 0.66 |
| *Fagus sylvatica* | 777.8 | 63.1 | 0.10 |
| *Quercus ilex* | 462.1 | 40.0 | 0.75 |
| *Pinus halepensis* | **140.7** | **7.2** | 0.68 |
| *Pinus nigra* | 656.1 | 57.4 | 0.57 |
| *Castanea sativa* | 584.1 | 49.0 | **0.61** |

## 4 Discussion

In this study, we applied the 3D-CMCC-FEM spatial explicitly over a Mediterranean region characterized by elevated bio-geographical and climatological heterogeneity. Initializing the forest structure, combining ground data from the NFI and the RS-based wall-to-wall basal area map, allows having a more realistic initial state of the carbon pools and forest structure, reducing, thus, uncertainties related to spin-up procedures which often translate into significant differences in the carbon fluxes (Carvalhais et al., 2008, 2010; Massoud et al., 2019; Lindeskog et al., 2021). The calibration of the allometric equations, to which the model has shown to be sensitive (Collalti et al., 2019), sets an additional constraint on the modeled initial growing biomass, which in turn influences the simulated GPP *via* the amount of sapwood. By means of the use of the basal area map to build the forest characteristics at the beginning of simulation, the model is able to retain part of the spatial information of the initial aboveground biomass and average forests structure, contributing, for instance, to the very satisfactorily spatial correlations of the modeled annual GPP with 2 out of 3 RS-based datasets. While aggregating basal area data at the 1 km resolution might reduce random uncertainties in basal area, additional uncertainty might stem from the tree density data, a data which is less constrained. Yet, performed tests (not shown here) indicate no significant sensitivity of the simulated GPP to stand density.

Overall, 3D-CMCC-FEM performances in simulating GPP when compared to other large scale and independent RS-based data, are shown to be generally satisfactory at both spatial as also temporal scales. In particular in the summer period across the seasons 3D-CMCC-FEM GPP show satisfactorily correlations and low relative differences and RMSE values when most of the vegetation activity takes place and it is thus the most robust signal across model and RS-based datasets. Yet, the comparative analyzed pinpointed some important challenges in applying the 3D-CMCC-FEM in Mediterranean areas and highlighted sources of uncertainties explaining the residual 3D-CMCC-FEM and RS-based data differences across RS-based datasets, which are grouped as follows:

### 4.1 3D-CMCC-FEM GPP vs. RS-based GPP

The 3D-CMCC-FEM GPP is close to the GOSIF and CFIX GPP estimates, while a general systematic positive overestimation emerge when comparing the 3D-CMCC-FEM GPP to FLUXCOM GPP (although for not all species, Figure 9). The 3D-CMCC-FEM and RS-based data differences resulting from the analyses are here discussed in relation to the different nature of the RS-based datasets used and their underlying algorithms and adopted approaches. FLUXCOM is an up-scaled product of the local EC-tower GPP estimates which is often used as benchmark RS-based dataset in model evaluation analyses (e.g. Byrne et al., 2018). Part of its worldwide application relies on the capability to sample in the entire climate-vegetation space. However, some particular areas at the rear edge of the Mediterranean forests might be not covered by this product (Jung et al., 2020). In addition, FLUXCOM operates also at coarse spatial resolution (Zheng et al., 2020; Zhang et al., 2022). Compared to GOSIF and CFIX, the spatial, seasonal and interannual variability in FLUXCOM relies on the RS-based dataset alone, without including any climatic drivers. Unfortunately, climate has been shown to play a significant role in the local, regional and global GPP. The lack of climatic drivers in FLUXCOM, might, thus, partly explain the systematic differences observed when compared to the 3D- CMCC-FEM GPP, which, at the opposite, showed to be sensitive to climate (Mahnken et al., 2022). However, the noticeable differences found, although some good correlations for some species, between the 3D-CMCC-FEM GPP and FLUXCOM, is common to other process-based and dynamic vegetation models comparative studies (Li & Xiao, 2019b; Jung et al., 2020; Zhang & Ye, 2021), and recent studies found that the GPP may be underestimated in the FLUXCOM- GPP in temperate areas (e.g. Bacour et al., 2019; Norton et al., 2019; Wild et al., 2022).

The GOSIF and the CFIX datasets, which 3D-CMCC-FEM GPP seems to reply better than FLUXCOM, have the advantage of having a finer spatial resolution (∼5 and 1 km versus ∼ 8 km in FLUXCOM) and, thus, contain more information, with the GOSIF having a globally higher continuous coverage *via* the original SIF data, which is the proxy of the photosynthetic activity at canopy scales (Sun et al., 2017; Li & Xiao, 2019a). CFIX is instead driven by a Light Use Efficiency model driven by NDVI, which is a vegetation index more suitable to investigate plant greenness rather than purely photosynthetic activities, as highlighted by Camps-Valls et al. (2021) and Walther et al. (2019). This inherent characteristic for NDVI might explain the lack of spatial correlation but also lower spatial correlations when compared to FLUXCOM and GOSIF. In addition, the Light Use Efficiency models are known to not saturate at increasing solar radiation, but that would lead CFIX having higher GPP values than 3D-CMCC-FEM GPP, which uses the biochemical model of Farquhar et al. (1980). However, while in some species CFIX has higher GPP values than 3D-CMCC-FEM GPP (e.g. *F. sylvatica* and *Q. Ilex*) in other species values are lower (*P. halepensis* and *Q. cerris*)(Figure 9). Differences between 3D- CMCC-FEM and CFIX may thus more probably rely on different models’ parameterization and not just on the different approach used to simulate photosynthesis. Indeed, as outlined in other studies carried at global scales there is a higher uncertainties in interannual variability even across different RS-based datasets, (Butterfield et al., 2020; Zhang & Ye, 2021) and even in its correct (of GPP) calculation across different models (Dunkl et al., 2023).

The differences in standard deviation between the 3D-CMCC-FEM and the RS-based GPP (see Figure S1), despite high in absolute terms, are still comparable to the range shown for instance in Zhang & Ye (2021) and Zheng et al. (2020). Similarly, the partial lack of agreement in the winter (but this is worth also for the spring and autumn) months, that we found in this study, it has been observed also in other RS-based data and process-based models comparative analyses (Zhang and Ye 2020). However, this it is not surprising, at least for deciduous species, given that 3D- CMCC-FEM simulates dominant vegetation only and no photosynthesis when there are no leaves on at the opposite to RS-based data which may account for underneath vegetation and this might explain the better correlations with evergreen species than for the deciduous for some RD-based datasets. Spring and autumn also depend on the spatial and temporal variability of bud breaks and leaf falls which control photosynthetic activity during and across the years. However, mismatches and asynchronies for the beginning and the end of the growing season largely vary between species and RS-based datasets considered as also found in the literature for other models (Peano et al., 2019, 2021). In addition, the RS-based datasets capture the fluxes embedding the entire sub-grid variability, including the contribution of the vegetation not simulated by the model (e.g. crops, *maquis* and understory which in any case comprise about one fifth of the entire vegetation area only), in particular in areas where the forest cover is indeed low because of low stand density, such as in the plain areas of the region under study or in degraded areas. Spatial scale mismatch between data and models may explain, thus, likely part of the low performances in some areas of the region. Zhao and Zu (2022) showed how GOSIF has apparently a similar interannual variability and trends when compared to the TRENDY simulations (an ensemble of simulations from Dynamic Global Vegetation Models; Sitch et al., 2015), as similarly as we found here using a process-based, stand-level, forest model.

On the other hand, some have raised doubts and concerns on the robustness of the relationship between GPP and remotely sensed sun-induced fluorescence (SIF)(Wohlfahrt et al., 2018; Bacour et al., 2019; Chen et al., 2021) showing that the SIF might overestimate the GPP values at least in temperate broadleaved forests (Qiu et al., 2020). However, to our knowledge, there are no specific indication that this retains also for Mediterranean forests. Yet the SIF data from RS remain one of the best proxies for photosynthesis over large areas to date (Sun et al., 2017; Li & Xiao, 2019a), making the GOSIF estimates potentially the more robust estimates compared to both CFIX and FLUXCOM.

Differences in the original spatial resolution of RS-based data or land cover used to drive the GOSIF and CFIX datasets, might additionally contribute to explain the 3D-CMCC-FEM GPP and RS-based datasets **–** but also between RS-based datasets **–** differences. Interestingly, when compared RS-based dataset vs. RS-based dataset GPP (e.g. GOSIF vs. FLUXCOM), and not just 3D-CMCC-FEM GPP vs. RS-based dataset, low correlations and discrepancies (both in the relative as in the absolute sense), although minor than against 3D-CMCC-FEM, across the results (see Table S3 and Table S4), both at the spatial as the temporal scale (including at the species level, see for example the large RMSE for all deciduous species in Table S4), have found. That was, however, expected given that RS-based methods are diagnostic and of similar nature tools rather than inherently prognostic (and with greater uncertainties) tools as potentially all models are. A recent large review on RS-based products intercomparison confirmed the large variability and different sensitivity to climatic factors between different approaches and criteria when compared at site-level (Sun et al., 2019).

### 4.2 3D-CMCC-FEM-related uncertainties

The 3D-CMCC-FEM model has been extensively evaluated at the site level all over Europe and evaluated against measurements of carbon fluxes from the FLUXNET database (Collalti et al., 2016, 2018; Marconi et al., 2017). The evaluation has been carried out in the past over a broad spectrum of climate, forest management, and species at stand level (e.g. Dalmonech et al., 2022; Testolin et al., 2023). In some recent model inter-comparison studies, the 3D-CMCC-FEM was shown to be able to simulate, among other things, C fluxes, e.g. GPP, and key structural variables, e.g. diameter, basal area, etc, with very good performance when compared to measured data and to other state-of-the-art forest models (Engel et al., 2020; Mahnken et al., 2022). However, when applying the 3D-CMCC-FEM at a different spatial scales model parameters, structural, and input-related uncertainties might amplify or even buffer the error in simulating GPP as observed for other models (e.g. Dalmonech et al., 2015; Zheng et al., 2022; Dunkl et al., 2023). In any case, the overall, 3D-CMCC-FEM GPP (∼600 - ∼2200 gC m^−2^ yr^−1^) is well in the bounds of the ∼600 - ∼2500 gC m^−2^ yr^−1^ GPP values described in Collalti and Prentice (2019) for temperate deciduous and coniferous forests in Europe.

The seasonal cycle of GPP describes the seasonal pattern of carbon gross assimilation by plants and the beginning and end of the growing season for the deciduous forests. For two out of the three datasets used here the 3D-CMCC-FEM may simulate a slightly earlier beginning of the growing season (e.g. for *Q. cerris*)(Figure S2). This might be partially attributable to the budburst parameterization, which is based on the Thermic sum and the growing degree days metric which is a trigger for the leaf flushing. Such a parameter is kept constant (as in many other models, see Peano et al., 2019; Peaucelle et al., 2019; Collalti et al., 2019) as a species-specific parameter, irrespective of any local climatic adaptation. The observed discrepancy in the beginning of the growing season, compared to the delay projected by the 3D-CMCC-FEM model, may be partly attributed also to the influence of undergrowth vegetation (such as grass or shrubs) on RS-based data. These understory plants frequently begin photosynthesis earlier, a phenomenon not accounted for by the 3D-CMCC-FEM model. The seasonality of the GPP in the evergreen species is instead apparently more shaped by the direct environmental effect on the photosynthetic process at a seasonal time scale (and less affected by underneath vegetation), suggesting that the 3D-CMCC-FEM GPP and RS-based datasets differences might be also attributable to some bias in the meteorological data used or for the other physiological parameters adopted or in the below-canopy vegetation. Indeed, model ecophysiological parameter values are derived from the literature, yet not calibrated for the specific sites or regions, in order to allow the 3D-CMCC-FEM general applicability as reported in Mahnken et al. (2022). In addition, other potential source of uncertainty, although minor, for the apparent mismatches in the beginning and in the end of the growing season between 3D-CMCC-FEM and RS-based datasets may stem from the different temporal resolutions, indeed, e.g. FLUXCOM is an 8-day product while 3D-CMCC-FEM is a daily one.

To capture the summer GPP sensitivity to hydrological variability, i.e. wetter or drier conditions, we use a simple GPP-SPEI diagnostics. The 3D-CMCC-FEM was first shown to be able to simulate comparably to the RS-based datasets the interannual variability (see Table 2) and then of the summer GPP and aridity signal over a large extent (see Figure 8). In large areas of the region, the 3D-CMCC-FEM and the RS-based datasets have similar responses in terms of GPP- SPEI5. Yet, in some areas, a too-strong model response to SPEI5 also emerges, indicating that the 3D-CMCC-FEM has a higher summer GPP-response to negative SPEI5 (i.e. drier conditions) compared to the RS-based datasets. In particular, results indicate a higher model sensitivity of GPP to SPEI for oaks-dominated forests clustered on higher elevations. Oak’s species are known to be an isohydric species, i.e. they weakly regulate stomatal openness under drought conditions and are more resilient to drought (Ripullone et al., 2020; Castellaneta et al., 2022), and it is possible that the high sensitivity of GPP to SPEI5 in the 3D-CMCC-FEM is a result either of a too strong control of VPD on stomatal conductance (Grossiord et al., 2020) or of a too strong soil moisture control on the photosynthesis (Crow et al., 2020; Fang et al., 2021). Summer period is the season where most of the annual vegetation activity takes place and reaches its maximum, and which variability is the most prominent feature in Mediterranean areas. As a matter of fact, in a previous study (see Collalti et al., 2016) was shown how, at site level in beech and coniferous forests, the 3D-CMCC-FEM was able to sufficiently simulate the sign of the year-to-year variability of the annual GPP anomalies, and in particular in years affected by important summer drought (i.e. 2003). Yet interannual variability is the most uncertain signal event when comparing different data and often considered as an ‘acid test’ in vegetation modeling (Keenan et al., 2012; Collalti et al., 2016; Dunkl et al., 2023).

However, stand-level studies showed how the 3D-CMCC-FEM model realistically simulates the response of GPP to atmospheric VPD and temperature (Mahnken et al., 2022). In the model soil hydrology is simulated *via* a single-layer bucket model and this might be another factor contributing to the stronger than observed modeled response, as it has been shown as overall that the majority of models with a soil bucket hydrology tend to limit GPP more than models with other soil conceptual e.g. multilayered schemes (Hanson et al., 2004). This higher sensitivity and the apparent stronger drop of modeled summer GPP might partially explain the higher standard deviation of the modeled signal compared to RS-based datasets (see Figure S1). One potential additional explanation to the modeled stronger GPP sensitivity to SPEI, would be the detected, although the narrow, earlier onset of the growing season in the model compared to the GOSIF dataset. In fact, leaf development not only drives how earlier or later the uptake of carbon from the atmosphere starts in the season, but also how earlier/later other processes, such as the leaf transpiration occurs in spring, potentially affecting the summer GPP response (Peano et al., 2019; Bastos et al., 2021; Chen et al., 2023).

### 4.3 Meteorological and soil data uncertainties

Low accuracy in the reconstructed meteorological forcing (see Bandhauer et al., 2022) might play an additional important role in studies that are conducted at high spatial resolution. The impact of the meteorological data used in data-driven and PBFMs was already shown to be important to accurately simulate GPP (e.g. Wu et al., 2017; F. Zhang et al., 2022; Y. Zhang & Ye, 2022). The 3D-CMCC-FEM was shown to be able to simulate GPP closer to observations at the site level in northern, central and southern Europe, where the downscaling and bias correction of the climate data was facilitated by the local geography (Collalti et al., 2018; Testolin et al., 2023).

Potential positive biases in the original E-OBS temperature dataset, as its accuracy relies also on the density of the termopluviometric stations and altitudinal cover, might translate into higher temperatures at higher elevations where, indeed, there are few stations as in southern Italy. A positive bias in temperature over areas where the temperature should be a limiting factor for vegetation, may partially explains the high modeled GPP values in beech forests leading to an anticipated onset or at thermic stress during the summer. Beech in the Italian Apennine Mountain regions represents the upper limit of forest vegetation and other studies (J. Fang & Lechowicz, 2006; Marchand et al., 2023) showed that the most limiting factor for the presence and growth of the beech in Europe is, indeed, temperature. The simulated values of GPP in beech-dominated forests are, on average higher than the three reference dataset, with simulated values in some areas larger than 2000 gCm^−2^yr^−1^. However these values are overestimated as observed average annual values of GPP in a beech forest in central Italy, a monitoring site with fluxes measured by means of the eddy covariance technique, are in the order of 1600 gC m^−2^yr^−1^ with range between ∼550 and 2500 gC m^−2^ yr^−1^ from Reyer et al. (2020). When applied in the same beech forest, the 3D-CMCC-FEM was shown to be able to fall within the observed range of GPP (Mahnken et al., 2022), indicating how the source of bias might be indeed attributable to uncertainties stemming from the up-scaling and affecting mostly the medium to higher elevation areas.

Simulated values for the deciduous oaks are indeed more comparable to EC-estimates in a site in central Italy (Roccarespampani, www.fluxnet.com, Tedeschi et al., 2006), with annual values of the order of 1600 gC m^−2^ yr^−1^for model and observations. However, a west-east gradient in the model-data difference is apparent at the regional level when comparing the 3D-CMCC-FEM to CFIX and GOSIF. In the former case, differences correlate with elevation and temperature, very likely a results of the lack of accuracy in the E-OBS dataset. Spatial differences in the 3D- CMCC-FEM GPP emerge in this study, which cannot be explained only by uncertainties in the modeling setup alone. The amount of water available for plants is determined in the 3D-CMCC-FEM by the balance between the precipitation (the inflow) and the Evapotranspiration (the outflow) as well as the soil characteristics, such as soil depth and texture. The water availability and the soil texture translate into the matric potential, which directly affects the stomatal regulation, leaf transpiration and the photosynthesis. When the model is applied spatially in a Mediterranean area with orographic and geographical complexity, the capability of the climate to retain the spatial and temporal variability of precipitation might have important effects in modulating the modeled spatial and temporal GPP variability (see Y. Zhang & Ye, 2022), and this will be the object of further investigations.

The last source of uncertainties in the 3D-CMCC-FEM GPP and RS-based GPP might stem from the soil characteristics and, in particular, the soil depth. The dataset used in the analyses as a proxy of depth available to root does not show any sensible spatial variability, e.g. with elevation, which is not realistic considering the topography and plasticity of plants roots (Fan et al., 2017; Stocker et al., 2023). The 3D-CMCC-FEM at this scale of application would benefit from coupling with more advanced hydrological scheme or a soil depth parameterization, to better couple the vegetation with the effective soil water availability (Kollet & Maxwell, 2008; Niu et al., 2011) or from soil moisture assimilation from remote sensing products (Kumar et al., 2014; Crow et al., 2020).

### 4.4 Limitations and further considerations

In this study we did not consider the land use change and the model does not simulate any spatial interaction from neighboring grid cells, not allowing for example simulating forest expansion. We additionally aggregated the forest species according to macro classes, hence neglecting intra- specific differences, if any. However, some compromises between input data requirements and time for model runs was needed to keep simulations and calculations feasible. In particular, at the spatial resolution of the study, we were more interested in capturing trends and spatial variability in GPP than the finest sub-grid variability within the forest ecosystem.

No Nitrogen cycle is included, however, the precipitation is commonly identified as the main limiting factor for photosynthesis in Mediterranean areas (Keenan et al., 2009; Flexas et al., 2014; Fyllas et al., 2017).

## 5 Conclusions and outlook

PBFMs offer a complementary tool to ground forest inventory networks and satellite-based records in monitoring carbon sequestration forest growth and the climate change impacts on forests, on many other variables which are otherwise difficult to measure extensively or monitor continuously. However, any model’s reliability needs to be verified and tested even in high heterogeneity contexts and not only to site-level ones. With this aim, we used RS-based data to initialize, apply, and test for the first time the biogeochemical, biophysical, process–based model 3D-CMCC-FEM on a regular grid at high spatial resolution under typical Mediterranean climate. We compare the obtained gross primary production over a large area (∼80% of the Basilicata region) against a suite of different in nature independent RS-based data. In spite of the simplified initial forest setup and the underlined uncertainties, the 3D-CMCC-FEM was shown capable of capturing both the spatial and the temporal variability of the RS-based data, even at species-level. Further tests are needed, yet the very promising results open the possibility of using the PBFMs to investigate the spatio-temporal dynamics of forest growth over larger spatial scales and under drought conditions and future climate scenarios, shaped by the spatial climatic and ecological heterogeneity such as in the Mediterranean areas.

## Acknowledgments

The work was carried out in the framework of the project ‘Advanced EO Technologies for studying Climate Change impacts on the environment – OT4CLIMA’ (D.D. 2261 - 6.9.2018, PON R&I 2014–2020 and FSC). This work has been partially supported by MIUR Project (PRIN 2020) “Unraveling interactions between WATER and carbon cycles during drought and their impact on water resources and forest and grassland ecosySTEMs in the Mediterranean climate (WATERSTEM)”(Project number: 20202WF53Z), “WAFER” at CNR (Consiglio Nazionale delle Ricerche) and D.D., E.V., A.C. were supported by PRIN 2020 (cod 2020E52THS) - Research Projects of National Relevance funded by the Italian Ministry of University and Research entitled: “Multi-scale observations to predict Forest response to pollution and climate change” (MULTIFOR, project number 2020E52THS). D.D., E.V., A.C. acknowledge also funding by the project OptForEU H2020 research and innovation programme under grant agreement No. 101060554. We also acknowledge the project funded under the National Recovery and Resilience Plan (NRRP), Mission 4 Component 2 Investment 1.4 - Call for tender No. 3138 of 16 December 2021, rectified by Decree n.3175 of 18 December 2021 of Italian Ministry of University and Research funded by the European Union – NextGenerationEU under award Number: Project code CN_00000033, Concession Decree No. 1034 of 17 June 2022 adopted by the Italian Ministry of University and Research, CUP B83C22002930006, Project title “National Biodiversity Future Centre - NBFC”. We acknowledge the E-OBS dataset from the EU-FP6 project UERRA (https://www.uerra.eu) and the Copernicus Climate Change Service, and the data providers in the ECA&D project (https://www.ecad.eu)”. The FLUXCOM products were obtained from the Data Portal at https://www.fluxcom.org/. D.D. thanks C. Trotta for valuable discussions about tree allometric relationships and thanks M. Willeit and E. Grieco for providing comments on a previous draft of the manuscript.

## Author contribution

**Daniela Dalmonech**: Conceptualization, Data curation, Formal analysis, Investigation, Methodology, Resources, Software, Validation, Visualization, Writing - original draft. **Elia Vangi, Gherardo Chirici,** and **Francesca Giannetti:** Resources, Writing - review & editing. **Jingfend Xiao**: Resources, Writing - review & editing; **Marta Chiesi** and **Luca Fibbi:** Resources, Writing - review & editing**. Gina Marano:** Software, Writing - review & editing **Christian Massari:** Writing - review & editing. **Angelo Nole**: Resources, Writing - review & editing. **Alessio Collalti**: Conceptualization, Formal analysis, Investigation, Methodology, Resources, Software, Validation, Visualization, Writing - review & editing.

## Data Availability Statement

The 3D-CMCC-FEM model code version 5.6 is publicly available under the GNU General Public Licence v3.0 (GPL) and can be found on the GitHub platform at: https://github.com/Forest-Modelling-Lab/3D-CMCC-FEM). All data and model executable, and scripts to perform analyses and figures presented in this work are provided open access in the Zenodo server (https://doi.org/10.5281/zenodo.8060401). Correspondence and requests for additional materials should be addressed to the corresponding author.

