## Supplementary Material for "Regional estimates of gross primary production applying the Process-Based Model 3D-CMCC-FEM vs. multiple Remote-Sensing datasets" (from another version of this preprint: version 1, 2023-06-14)

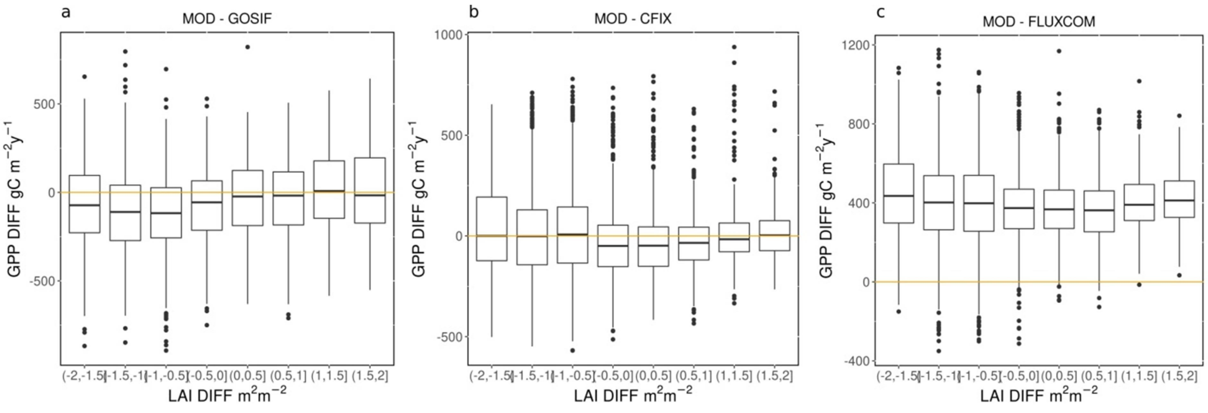


**Figure S1| Differences between modeled annual GPP and satellite based GPP estimates of a) GOSIF-GPP, b) FLUXCOM-GPP and c) CFIX-GPP, reported according to binned differences between multiyear-average of leaf area index (LAI) annual maximum between model and MODIS dataset (Product MCD15A3H.061, Myneni et al.2002,2015). The period of reference is 2005-2013.**

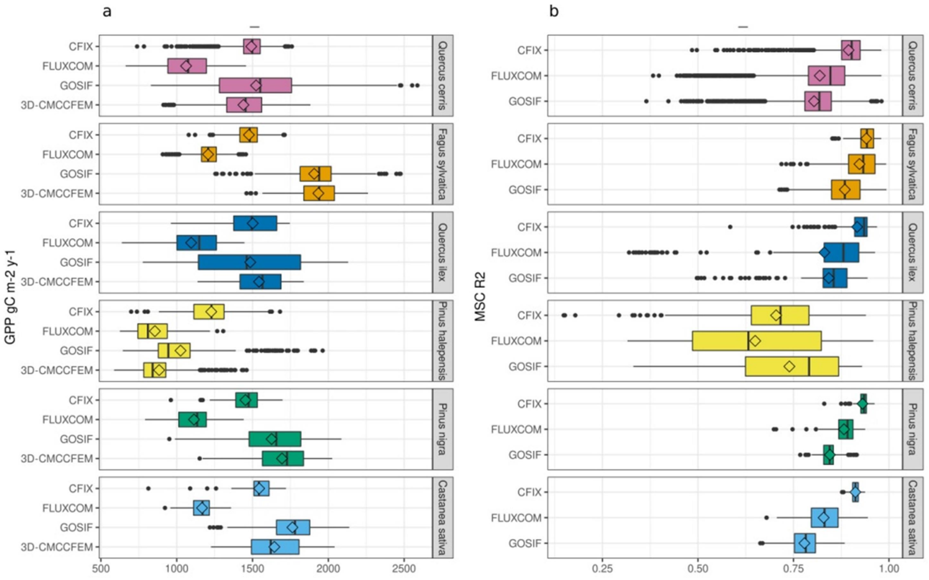

**Figure S2 | a) Box plot of the multiyear-average annual GPP data for model and the satellite-based datasets (GOSIF-GPP, CFIX-GPP and FLUXCOM-GPP). b) box plot of the $R^2$ of the GPP Mean Seasonal cycle between model and datasets. Analyses were performed for the years 2005-2013 and data aggregated according to the forest species.**

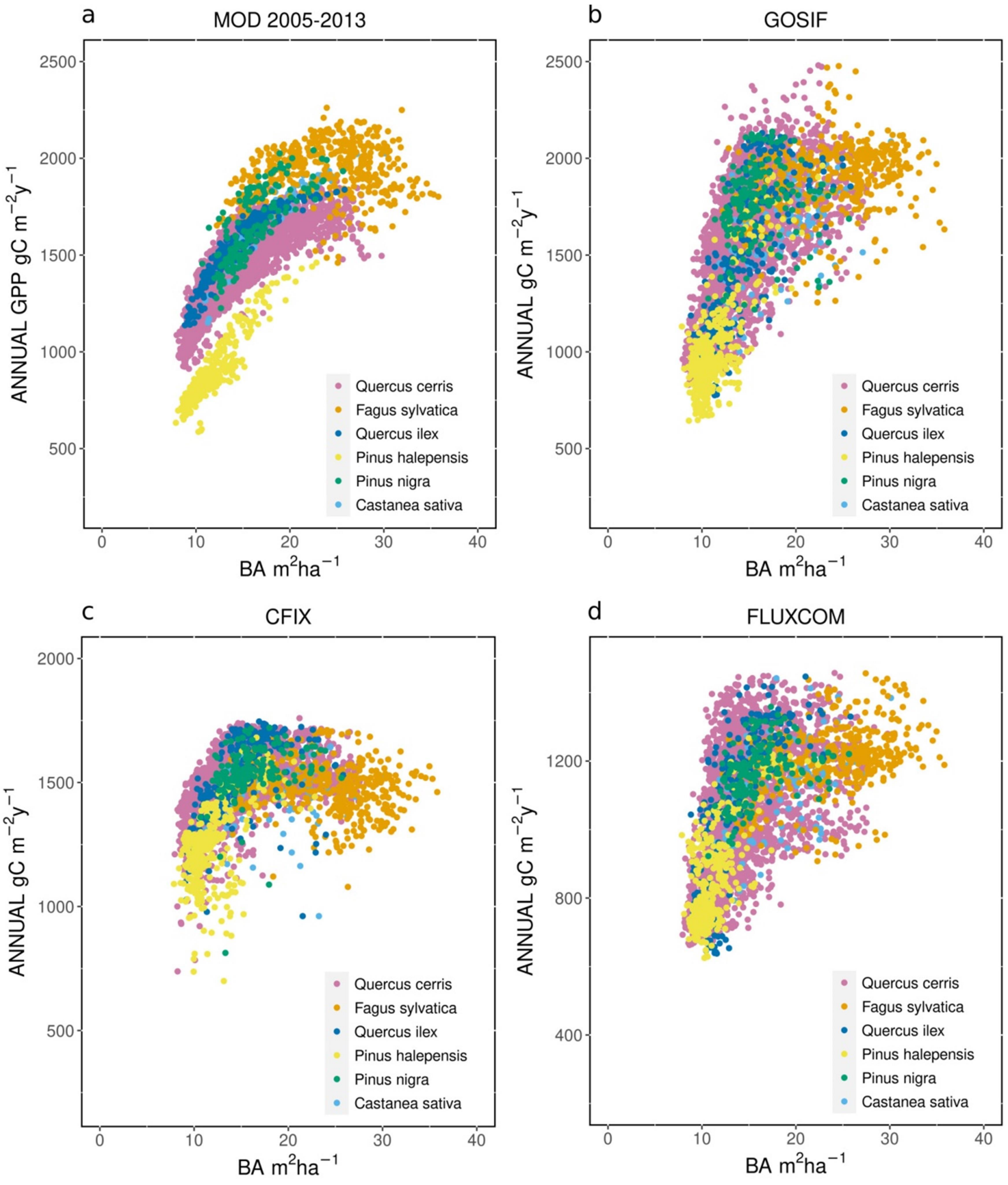


**Figure S3 | Annual GPP values over the period 2005-2013 for a) 3D-CMCC-FEM model, b) GOSIF-GPP, c) CFIX-GPP , d) FLUXCOM-GPP plotted versus the Basal Area (BA) values aggregated for each grid cells and dominant forest class at year 2005. Different dominant classes are reported in different colour, The plot is a descriptive diagnostic to**

**visually compare the GPP~BA pattern in model and datasets, for this reason the y-axes do not show the same range.**

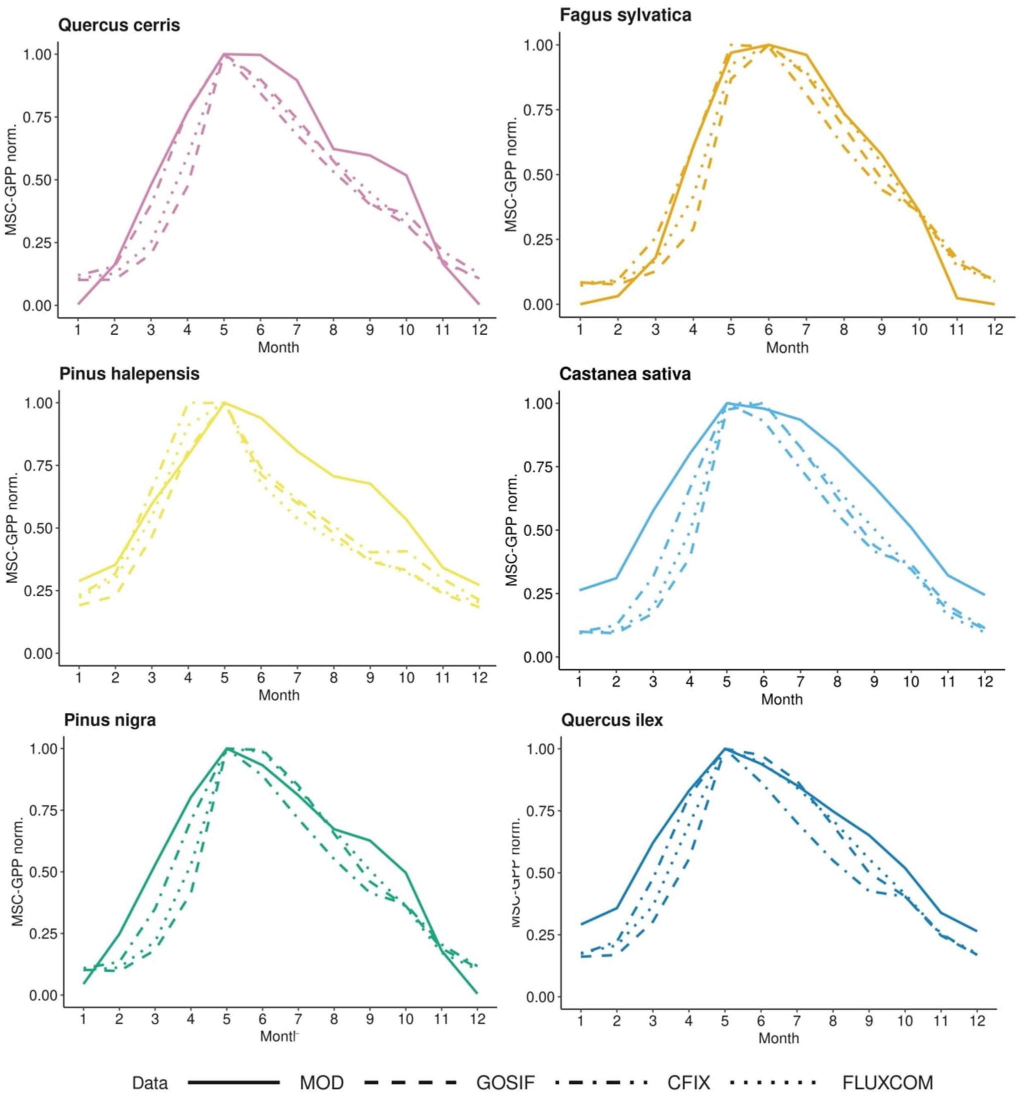


**Figure S4 | Normalized mean seasonal cycle of GPP for model and reference datasets plotted according to forest class. The mean seasonal cycle is computed for the period 2005-2013.**

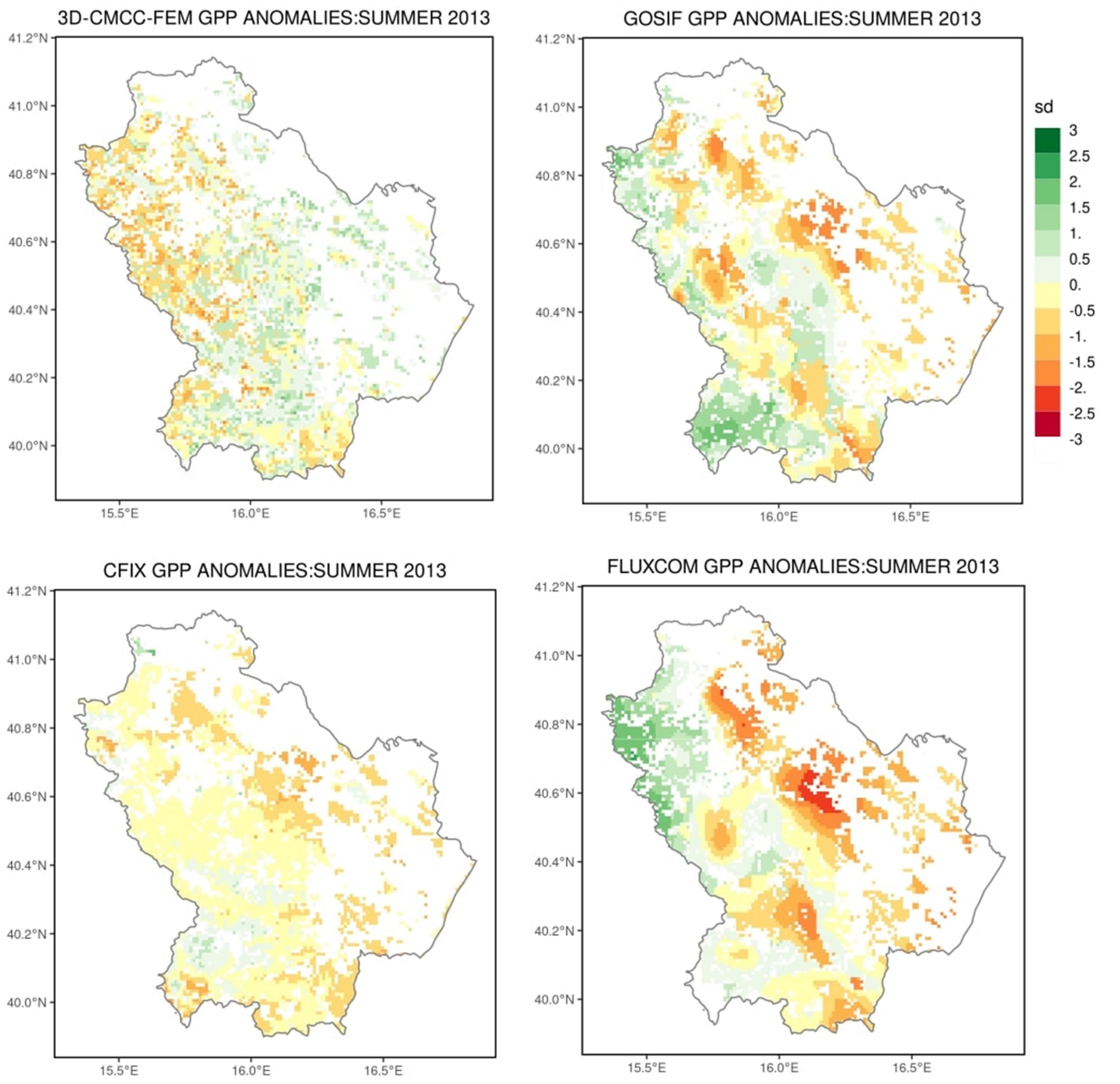


**Figure S5 | Standardized anomalies for the summer season (June-July-August) were calculated by removing long-term linear trend from the data and then dividing by the corresponding standard deviation of the GPP. This has been performed for each grid cell and for the three reference datasets and the model. Only spatial anomaly for the summer 2013 is shown. Please note the different original spatial resolution across datasets: CFIX-GPP, 1 km: GOSIF-GPP, 5 km; FLUXCOM-GPP, ~ 8km.**

| FOREST CLASSES | ASSOCIATED AVERAGE AGE [y] |
|---|---|
| Coppice | 30 |
| Old Coppice | 45 |
| 'Transition' forest | 50 |
| Very Young Timber | 20 |
| Young Timber | 40 |
| Old Timber | 70 |
| Very old Timber | 90 |

**Table S1 | Correspondence between the stand development stage in the regional forest thematic map of Basilicata and the average age. The class 'Transition forest' indicates stands under transition from coppice to high forest.**

Because of the lack of extensive site level measurement or estimates, the choice of the age class was not well constrained. However, sensitivity analyses performed on the model showed the low importance of the initial age to the carbon variable output (Collalti et al., 2019).
